# Conserved Filovirus Proteins as Targets of Broad-Spectrum Antivirals

**DOI:** 10.1101/2025.09.26.678902

**Authors:** Marcus Tullius Scotti, Enes Kelestemur, Holli-Joi Martin, Brandon Novy, Cleber C. Melo-Filho, Renata Priscila Barros de Menezes, Chonny Herrera-Acevedo, Alexander Tropsha, Eugene N. Muratov

**Affiliations:** Laboratory for Molecular Modeling, Division of Chemical Biology and Medicinal Chemistry, UNC Eshelman School of Pharmacy, University of North Carolina, Chapel Hill, NC, 27599, USA; Department of Chemistry, Center for Exact and Natural Sciences (CCEN), Federal University of Paraiba (UFPB), João Pessoa 58051-900, Brazil; Postgraduate Program in Natural and Synthetic Bioactive Products (PgPNSB), Federal University of Paraiba (UFPB), João Pessoa 58051-900, Brazil; Department of Chemical Engineering, Universidad ECCI, Bogotá, Distrito Capital 111311, Colombia

**Author notes:** These authors contributed equally (Marcus Tullius Scotti, Enes Kelestemur).

**Keywords:** Filoviridae, Ebola, Marburg, Broad Spectrum Antiviral, Protein Homology

## Abstract

Filoviruses are enveloped, non-segmented, negative-strand RNA viruses belonging to the *Filoviridae* family, which includes five genera: *Ebolavirus*, *Marburgvirus*, *Cuevavirus, Striavirus*, and *Thamnovirus*. Members of this family cause severe and, often, fatal hemorrhagic fevers in humans and non-human primates, with high mortality rates. To date, only two filoviruses, Ebola virus (EBOV) and Marburg virus (MARV), are known to infect humans and are listed as priority pathogens by the World Health Organization due to their potential for re-emergence and the current lack of effective vaccines and antiviral treatments. In this study, we identify and characterize conserved binding sites within key filoviral proteins to support the development of broad-spectrum, direct-acting antiviral agents. We validated the significance of these conserved regions for drug discovery using existing experimental data. Our analysis revealed notably high sequence similarity among proteins from filoviruses capable of infecting humans (EBOV, TAFV, BDBV, SUDV, MARV, and RAVV) compared to those from non-zoonotic species, with the highest conservation observed in the L and VP40 proteins—both critical for viral genome transcription and replication. Furthermore, we compiled and analyzed available experimental data on known antiviral compounds targeting these proteins, identifying several agents with cross-filovirus activity, including Galidesivir, Remdesivir, and Favipiravir. The integrated approach described here—combining sequence and structural conservation analysis with chemical structure and antiviral activity data—demonstrates a strategy that could be extended to the development of broad-spectrum therapeutics across multiple viral families.

**HIGHLIGHTS:**

- Conserved filovirus sites targeted for broad-spectrum antivirals.
- Structural modeling identifies key antiviral binding sites.
- Viral internal proteins are crucial targets for inhibition.
- Remdesivir validates conserved polymerase as a druggable target.
- Study highlights need for pan-filovirus drug screening

**TOC GRAPHIC:** 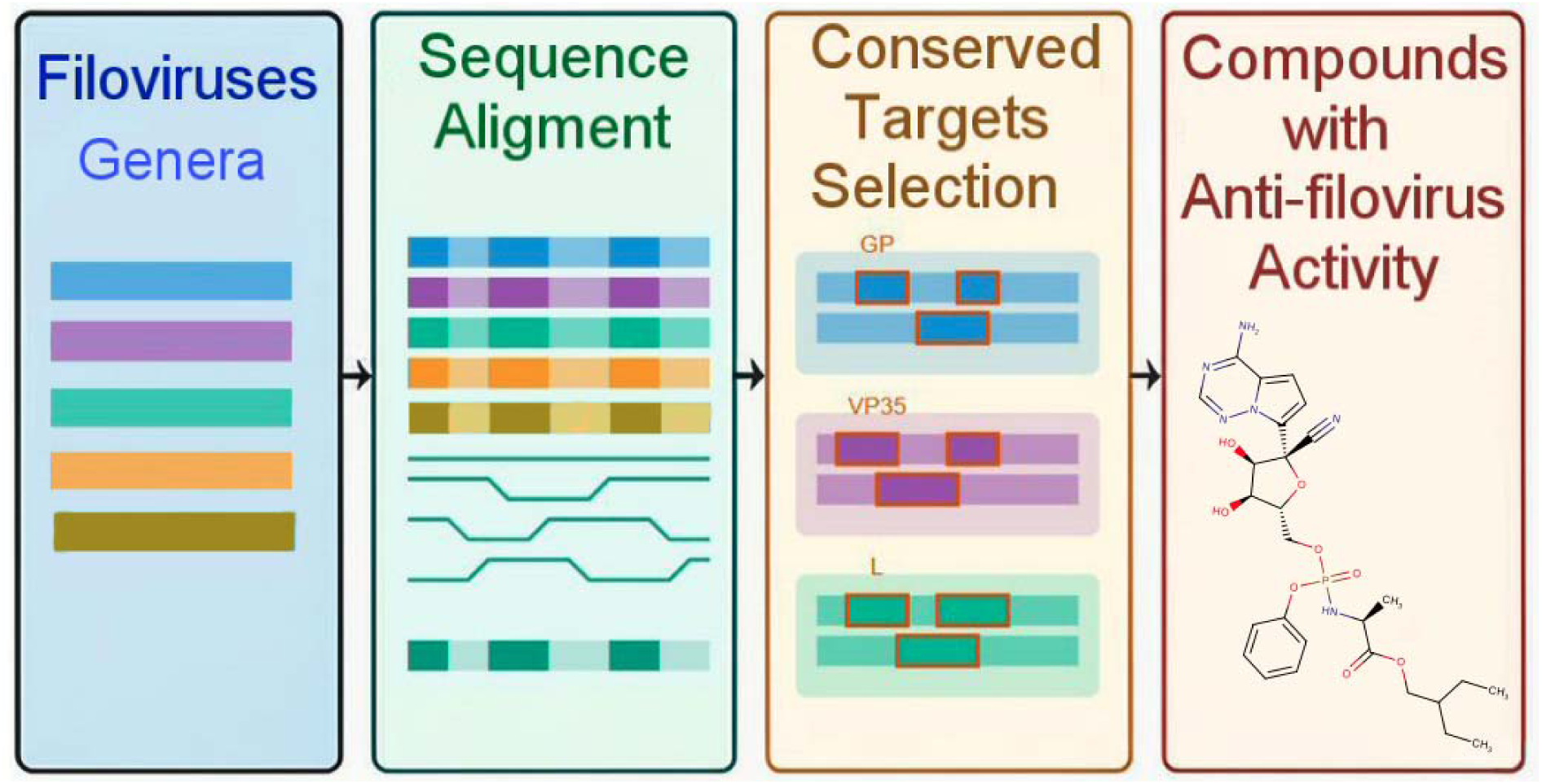

## 1. Introduction

Viruses of the *Filoviridae* family cause severe to fatal hemorrhagic fevers, with high mortality rates in humans and non-human primates [1,2]. This family comprises five recognized genera (*Ebolavirus*, *Marburgvirus*, *Cuevavirus*, *Striavirus*, and *Thamnovirus*), with an ongoing proposal to establish a sixth genus, *Dianlovirus* [3]. Filoviruses are enveloped, non-segmented, negative-strand RNA viruses that encode seven structural proteins: nucleoprotein (NP), viral proteins 24 (VP24), 30 (VP30), 35 (VP35), 40 (VP40), glycoprotein (GP), and the large (L) protein [2–5]. Among these, the L protein is particularly critical as it is the sole viral enzyme, harboring essential domains such as the methyltransferase (MTase) and the RNA-dependent RNA polymerase (RdRp). The RdRp domain governs viral RNA replication and transcription, positioning the L protein as a prime target for antiviral drug development [6–8].

Currently, only two filoviruses, Ebola virus (EBOV) and Marburg virus (MARV), are known to infect humans. Both viruses are classified as priority pathogens by the World Health Organization due to their potential for re-emergence and the absence of widely effective vaccines or antivirals. In various regions of Africa, Ebola virus outbreaks have led to devastating epidemics, with reported mortality rates ranging from 25% to over 90%, depending on the viral strain and healthcare infrastructure [5,9,10]. Additionally, a recent outbreak of Marburg virus in the Republic of Rwanda, ongoing since September 2024, has resulted in the largest number of infections recorded since 2005 [11]. More outbreaks of filoviruses are predicted in the near future [12].

In this study, we systematically identify and investigate conserved regions within key filoviral proteins to assess their potential as targets for broad-spectrum, direct-acting antiviral agents. Our approach involved analyzing the sequence conservation of all available filovirus proteins and evaluating binding site similarities for those with resolved three-dimensional (3D) structures deposited in the Protein Data Bank (PDB). We also compiled and reviewed relevant data on antiviral compounds tested against these targets. Building on our prior work on conservation-based antiviral discovery in coronaviruses [13], we highlight the utility of this integrated approach to accelerate the development of broad-spectrum therapeutics in preparation for future filovirus outbreaks.

## 2. Materials and Methods

### 2.1 Protein selection and collection

We identified amino acid sequences for twelve different species from the Filoviridae family and downloaded them from UniProt [14,15]—*Ebolavirus*: BOMV, TAFV, BDBV, RESTV, SUDV, and EBOV; *Marburgviruses*: MARV and RAVV; *Dianlovirus*: MLAV; *Cuevavirus*: LLOV; *Striavirus*: XILV; and *Thamnovirus*: HUJV (Table 1).

**Table 1:** Information about the viruses of the Filoviridae family analyzed in this study.

| Abbreviation | Genus | Species | Virus | Infects Humans |
| --- | --- | --- | --- | --- |
| **EBOV** | *Ebolavirus* | *Zaire ebolavirus* | Zaire virus | Yes |
| **SUDV** | *Ebolavirus* | *Sudan ebolavirus* | Sudan virus | Yes |
| **TAFV** | *Ebolavirus* | *Tai Forest ebolavirus* | Tai Forest virus | Yes |
| **BDBV** | *Ebolavirus* | *Bundibugyo ebolavirus* | Bundibugyo virus | Yes |
| **RESTV** | *Ebolavirus* | *Reston ebolavirus* | Reston virus | No |
| **BOMV** | *Ebolavirus* | *Bombali ebolavirus* | Bombali virus | No |
| **MARV** | *Marburgvirus* | *Marburg marburgvirus* | Marburg virus | Yes |
| **RAVV** | *Marburgvirus* | *Marburg marburgvirus* | Ravn virus | Yes |
| **MLAV** | *Dianlovirus* | *Mengla dianlovirus* | Mengla virus | No |
| **LLOV** | *Cuevavirus* | *Lloviu cuevavirus* | Lloviu virus | No |
| **XILV** | *Striavirus* | *Xilang striavirus* | Xilang virus | No |
| **HUJV** | *Thamnovirus* | *Huangjiao thamnovirus* | Huangjiao virus | No |

### 2.2 Sequences Alignment

The Clustal W alignment tool [16] was utilized to assess the percentage of identity among the aligned sequences of each protein within the analyzed species of the *Filoviridae* family (table 1). Amino acid similarities, expressed as percentages across all sequences, were identified, and the resulting alignments were saved in FASTA format.

### 2.3 Exploring Protein Similarities through Alignment Visualization

To visualize and analyze these similarities, heat maps and dendrograms were generated using KNIME 5.1.2 [17]. The similarity percentages were obtained using Clustal W, which employs a progressive multiple sequence alignment algorithm based on global pairwise alignments and employed to construct a graphical representation. The workflow integrated “Heat map” and “Hierarchical clustering” nodes, with the latter configured to execute three-cluster hierarchical clustering using the Euclidean Distance Function.

The visualization of multiple sequence alignments, homology searches, and assessment of secondary structure components was carried out utilizing ENDscript (Available on: https://endscript.ibcp.fr) and ESPript (Available on: https://espript.ibcp.fr) [18,19]. These tools facilitate the rapid exploration of protein structures and the identification of conserved regions.

All alignments generated by Clustal W were used as input files in ALN format. Subsequently, three-dimensional 3D structures retrieved from the Protein Data Bank (PDB) were employed as references for depicting secondary structures. Specifically, the PDB entries 4ZTG for NP, 4IBC for VP35 [7], 6G95 for GP [20], 4LDB for VP40 [21], 5T3T for VP30 [22], and 4M0Q for VP24 [23] were selected. It is noteworthy that these proteins were chosen from *Zaire ebolavirus* (EBOV) due to EBOV having the highest case-fatality rate among filoviruses capable of infecting humans, being linked to nearly two-thirds of *Ebolavirus* outbreaks [24,25].

## 3. Results and discussion

### 3.1 Comparative Analysis of Primary Protein Sequences Across Twelve Filoviridae Species

Initially, the similarity between the primary sequences of the seven structural proteins from 12 species of *Filoviridae* was analyzed (Figure 1). The *Zaire ebolavirus* (EBOV) was used as a reference, given that it is the virus species that infects humans and has shown a higher rate of cases and mortality [24]. Figure 1 illustrates that among *Ebolavirus* genus species capable of infecting humans, such as TAFV, BDBV, and SUDV, there is a notable higher percentage of similarity with our reference.

**Figure 1.**
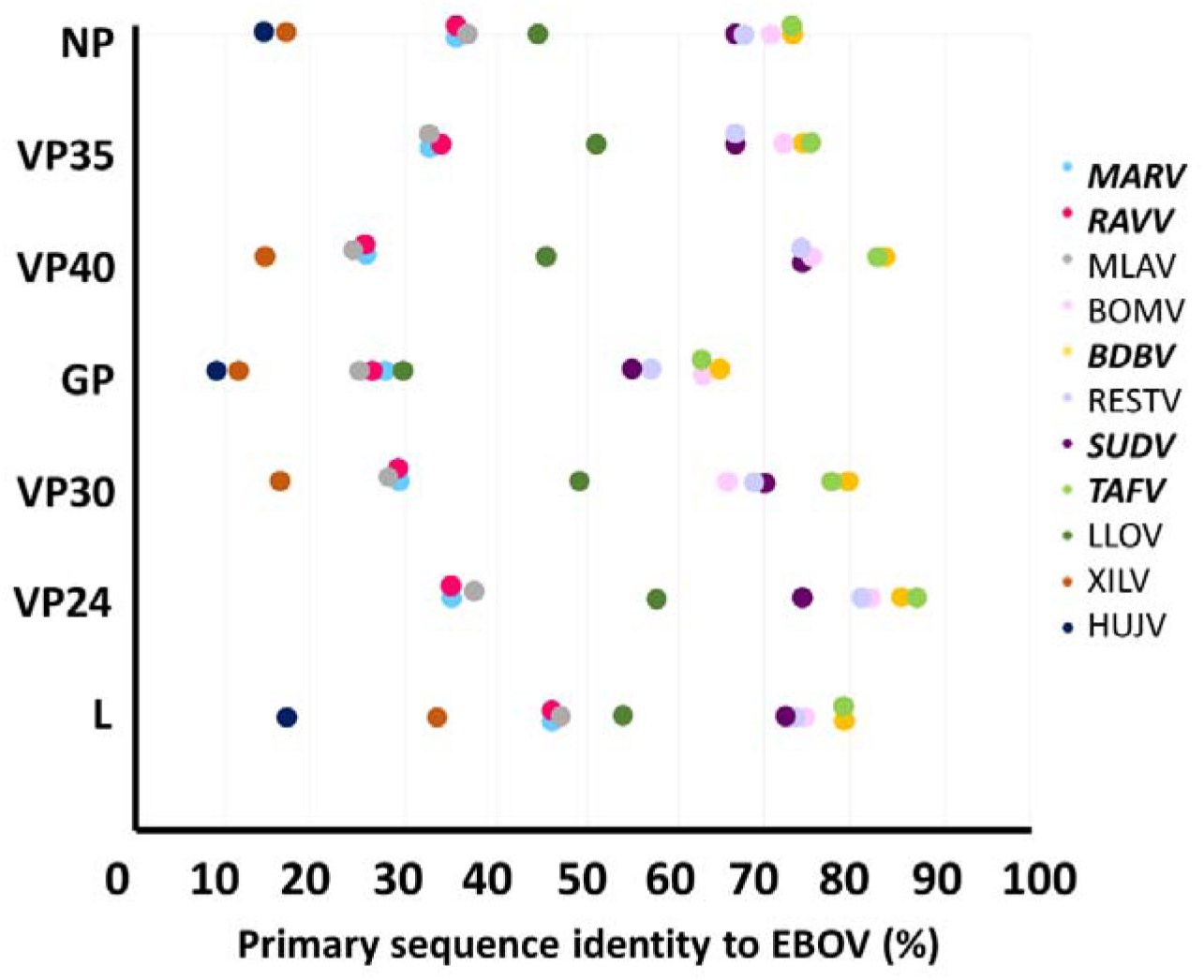
Primary sequence identity for structural proteins from various Filoviridae species, EBOV is used as reference. Bolded and italicized species are known to be infectious to humans.

BDBV and TAFV exhibit sequence identity values exceeding 72% across six of the seven structural proteins, with the exception of the glycoprotein (GP), which consistently shows lower conservation across all analyzed species. For all proteins assessed, TAFV and BDBV share approximately 10% higher identity with EBOV compared to SUDV. This higher degree of similarity may partially explain the comparable pathogenic potential observed between BDBV, EBOV, and SUDV, all of which have been linked to major disease outbreaks in Africa, whereas RESTV and TAFV, showing lower similarity, have not been associated with human epidemics [26].

In *Marburgvirus* species, MARV and RAVV, both capable of causing severe hemorrhagic fever, demonstrate markedly lower sequence similarity to members of other genera, with identity percentages generally below 40%. This sharp divergence highlights the evolutionary distance between *Marburgvirus* and *Ebolavirus* species, suggesting distinct mechanisms of pathogenesis and immune evasion. Interestingly, in EBOV, only the RNA-dependent RNA polymerase (RdRp) domain surpasses this threshold, reaching 47% identity, pointing to a relatively higher evolutionary pressure to conserve the enzymatic functions necessary for viral replication.

Figure 2 provides a visual summary of these patterns through heat maps illustrating identity percentages among *Filoviridae* species for six structural proteins. Within *Marburgvirus* species (MARV and RAVV), sequence conservation remains remarkably high, with identity values exceeding 90% across all proteins (Figures 2a, 2c–f), except for the GP (Figure 2b), which shows slightly lower conservation (∼77%). These findings suggest that despite their divergence from *Ebolavirus* species, MARV and RAVV maintain strong internal conservation, reflecting shared functional constraints essential for viral fitness. Furthermore, the distinct clustering of *Marburgvirus* and *Ebolavirus* species in the heat maps—highlighted by contrasting green and red regions—underscores the genetic and likely functional bifurcation between these two groups, which has important implications for the development of genus-specific versus pan-filovirus therapeutic strategies.

**Figure 2.**
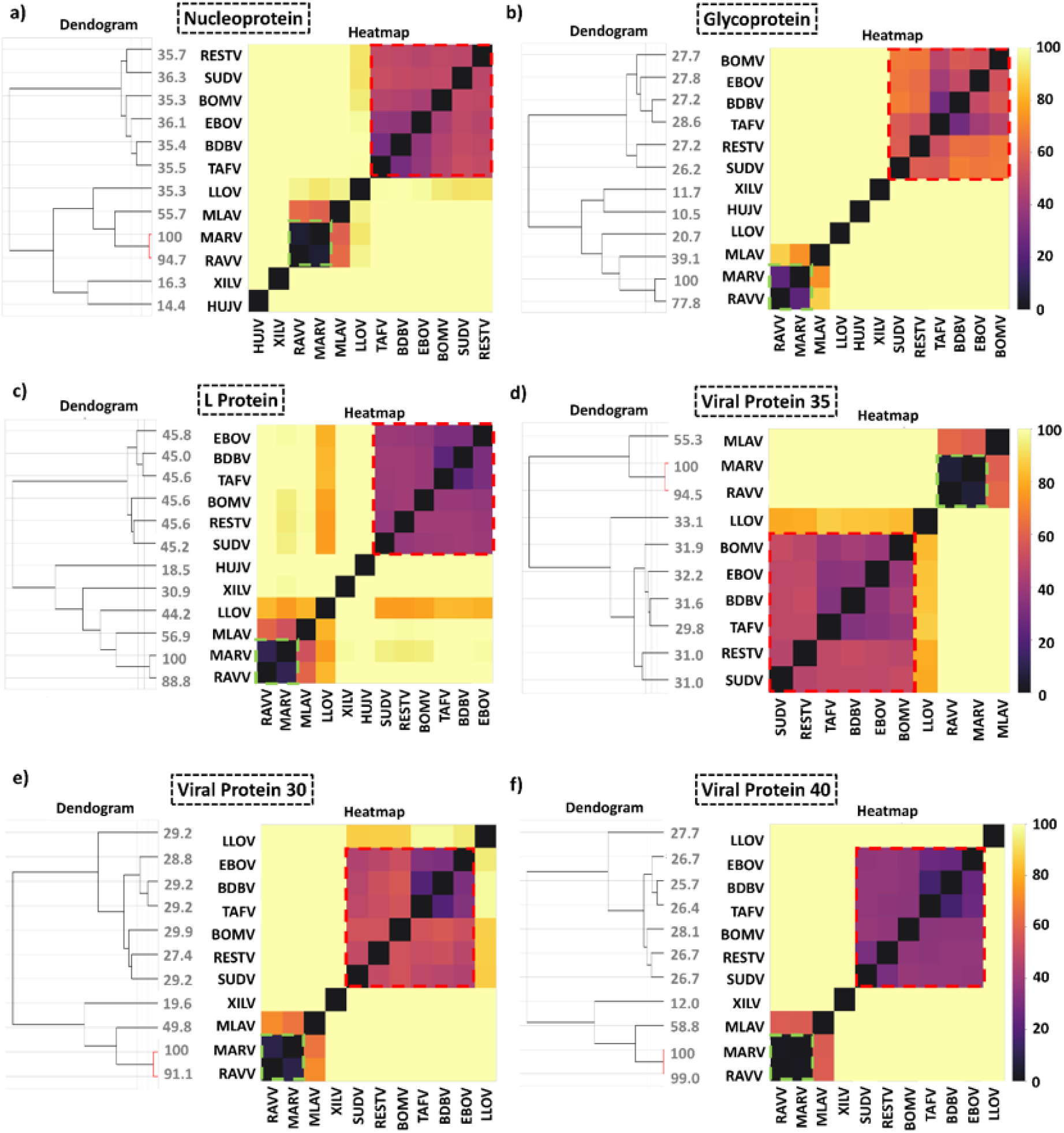
Primary sequence comparison of a) NP, b) GP, c) L, d) VP35, e) VP40, f) VP30 from different members of the Filoviridae family. The higher the numbers on the dendrograms – the higher the similarity of a given protein among the viral strains. Larger values indicate that sequences joined at a higher similarity level. The region highlighted in green corresponds to Marburg viruses and the region highlighted in red corresponds to Ebola viruses.

Similarly, the red regions in Figure 2 highlight the high degree of sequence conservation among *Ebolavirus* species relative to other members of the *Filoviridae* family. Supporting this observation, Baker et al. demonstrated through crystallographic analysis that the structures of BDBV and TAFV proteins closely resemble those of EBOV, underscoring the strong structural conservation within this genus [27].

Within *Ebolavirus* species, the L protein and VP40 protein display the highest levels of sequence similarity. Both proteins play critical roles in the viral life cycle: VP40 is a major structural component involved in virus assembly and budding, and the L protein, as part of the RNA-dependent RNA polymerase (RdRp) complex, is essential for viral genome replication and transcription. Notably, VP40 alone can efficiently drive the formation of virus-like particles (VLPs), a feature that highlights its pivotal role in viral egress. In contrast, VP30, which also interacts with the L protein as part of the RdRp complex, exhibits approximately 10% lower sequence identity compared to L and VP40 [28], suggesting a slightly greater tolerance for sequence variation without loss of function.

Interestingly, among the three species analyzed from the *Cuevavirus*, *Striavirus*, and *Thamnovirus* genera, LLOV (Lloviu virus) exhibited a higher percentage of sequence identity when compared with both *Marburgvirus* and *Ebolavirus* species. While the overall genome organization of *Filoviridae* members remains highly conserved—with similar gene order and protein functions—significant sequence divergence persists between the major genera, particularly *Ebolavirus*, *Cuevavirus*, and *Marburgvirus* [29]. These patterns of conservation and divergence reflect underlying evolutionary pressures and may influence host range, pathogenicity, and immune evasion strategies across filovirus species.

All these observations are summarized in Figure 3, where hierarchical clustering was performed using KNIME 5.2.1 based on the sequence identity of the 12 studied *Filoviridae* species. The analysis revealed three primary clusters. The first cluster comprises the *Marburgvirus* species, with the *Dianlovirus* member MLAV also grouped within this cluster. MLAV has previously been shown to be closely related to MARV among filoviruses [30]. Notably, *Dianlovirus* was isolated from *Rousettus* bats in China, and its genome shares high similarity with MARV strains found in African *Rousettus* bats, the known natural reservoir of Marburg virus [31]. This close relationship supports the hypothesis that similar ecological niches and host reservoirs contribute to the genetic conservation observed within this group.

**Figure 3.**
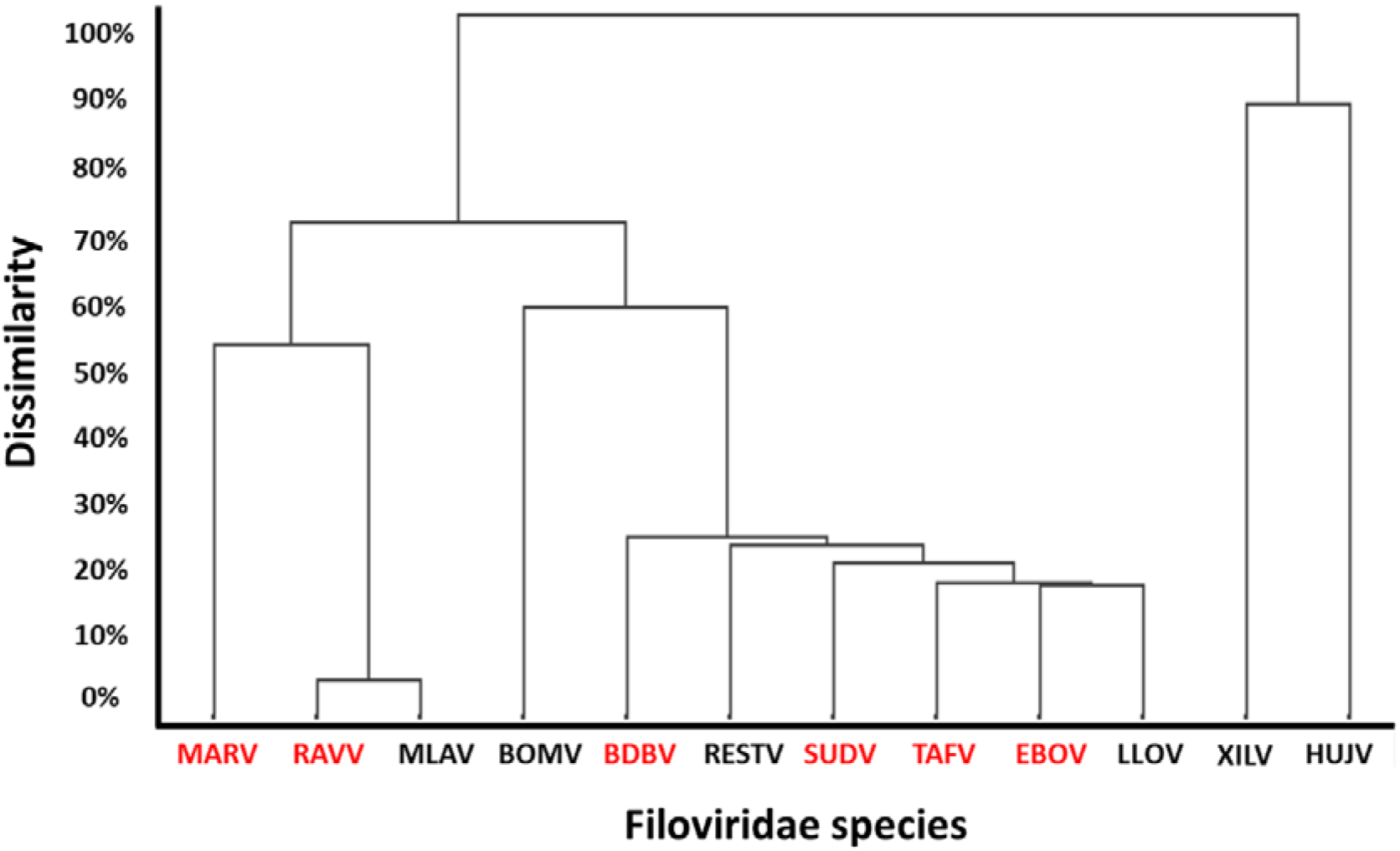
Hierarchical clustering of Filoviridae species, with human-infecting species highlighted in red.

The second cluster contains the six *Ebolavirus* species—BOMV, TAFV, BDBV, RESTV, SUDV, and EBOV—along with the *Cuevavirus* LLOV. Interestingly, LLOV clusters closer to *Zaire ebolavirus* (EBOV), indicating a relatively higher sequence similarity compared to other filoviruses. Within the *Ebolavirus* cluster, BOMV displays the highest degree of sequence divergence. This greater variability may explain its distinct biological properties, including a slower in vitro growth rate compared to EBOV and a markedly lower pathogenicity in humanized mouse models—findings consistent with earlier observations for RESTV, the only *Ebolavirus* species of Asian origin [25,32].

The third cluster comprises *Striavirus* (XILV) and *Thamnovirus* (HUJV), two filoviruses that, to date, have not been associated with mammalian hosts. Instead, they have been detected in actinopterygian fish species from Asia—specifically in frogfish (XILV) and filefish (HUJV) [33,34]. Notably, due to the incomplete genomic characterization of these viruses, sequences for several structural proteins were unavailable and thus were not included in the sequence alignment analysis. Together, these clustering patterns reinforce the evolutionary divergence among filovirus genera and provide valuable insights into their host adaptation and zoonotic potential.

### 3.2 Binding site similarity

To investigate the active site of the proteins, protein alignment was carried out, with all species of viruses which included six species of *Ebolavirus* EBOV, SUDV, TAFV, BDBV, RESTV, and BOMV, two *Marburgvirus* species: MARV and RAVV, one *Dianlovirus:* MLAV, one *Cuevavirus:* LLOV, one *Striavirus:* XILV and one *Thamnovirus*: HUJV. Proteins from the *Zaire ebolavirus* species, a member of the *Ebolavirus* genus, were used as a reference. This alignment aimed to analyze the conservation of amino acid residues in the active sites of the NP, VP24, VP30, VP35, VP40, GP, the L protein. The most conserved protein binding sites are shown in Figure 4.

**Figure 4.**
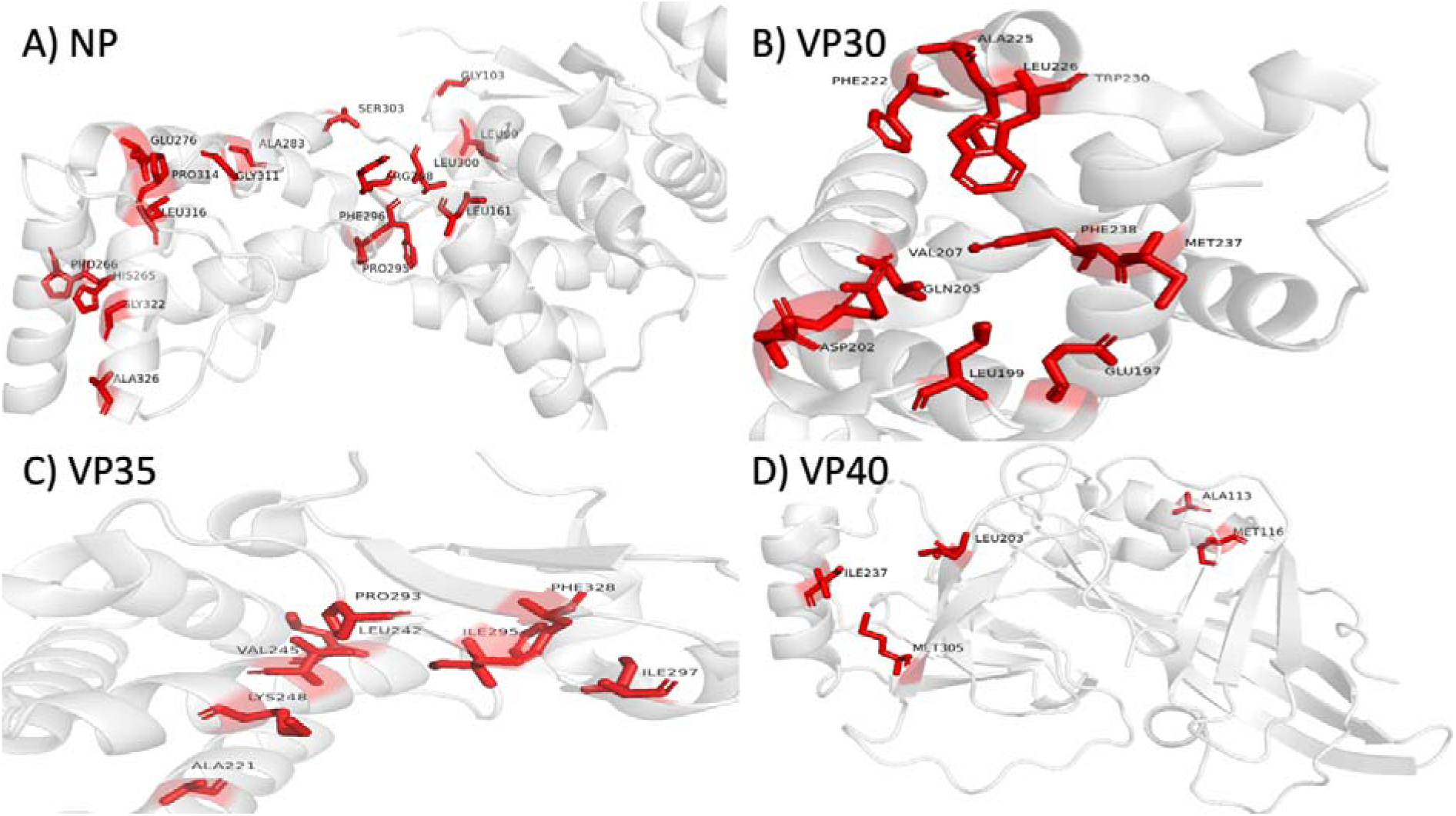
Binding site residues of conserved and semi-conserved NP (A), VP30 (B), VP35 (C), and VP40 (D) proteins in all homologous viruses listed in Table 1.

#### Viral Protein 35 (VP35)

Viral protein 35 (VP35) is a highly conserved and multifunctional protein across *Filoviridae* species, playing critical roles in viral replication and immune evasion. As a cofactor in the viral RNA-dependent RNA polymerase (RdRp) complex, VP35 is essential for facilitating viral genome replication and the transcription of negative-sense viral mRNA [35,36]. Beyond its role in replication, VP35 functions as a potent interferon (IFN) antagonist, effectively inhibiting host innate immune responses and promoting viral survival [36,37].

A key additional function of VP35 is its interaction with the nucleoprotein (NP). VP35 binds to the N-terminal domain of NP, preventing NP oligomerization and maintaining it in a monomeric form until it associates with viral RNA. This VP35–NP interaction is crucial for the proper assembly of the polymerase complex, with VP35 serving as a bridge between NP and the L polymerase, ultimately ensuring efficient viral RNA synthesis [36–38]. Through these complementary roles, VP35 emerges as a central player in both the replication cycle and immune evasion strategies of filoviruses.

Structurally, VP35 contains a well-defined active site pocket composed of approximately 20 amino acid residues arranged within α-helical and β-sheet subdomains. Key residues contributing to inhibitor binding within this pocket include Ala221, Arg225, Gln241, Leu242, Lys248, Lys251, Pro293, Ile295, Ile297, Asp302, and Phe328. A prior structural analysis by Brown et al. (2014) [28] identified Val245, Ile295, Phe328, Lys251, and Gln241 as critical residues essential for inhibitor interaction and enzymatic regulation.

Protein sequence alignment across multiple filoviruses revealed notable conservation within the VP35 active site. Specifically, residues Ala221, Leu242, Val245, Lys248, and Pro293 are fully conserved across all analyzed species, underscoring their fundamental role in maintaining VP35’s structural and functional integrity. Residues Ile295, Ile297, and Phe328 are semi-conserved, showing minor variations while preserving overall chemical properties important for structural stability. Conversely, residues Arg225, Gln241, Lys251, and Asp302 are semi-conserved only within the *Ebolavirus* (EBOV, SUDV, TAFV, RESTV, BOMV, BDBV) and *Cuevavirus* (LLOV) genera but are not conserved in *Marburgvirus* (MARV, RAVV) and *Dianlovirus* (MLAV), suggesting genus-specific structural adaptations.

These findings, illustrated in Figure S1, highlight the evolutionary constraints acting on VP35’s active site and reinforce its potential as a promising, broad-spectrum antiviral target, particularly for therapeutics aimed at *Ebolavirus* and *Cuevavirus* infections.

#### Viral Protein 40 (VP40)

Viral protein 40 (VP40) is a multifunctional structural protein critical for the life cycle of filoviruses. It exhibits remarkable conformational flexibility during infection, enabling the virus to adapt to different stages of assembly and egress. Initially, VP40 localizes at the host cell plasma membrane as a butterfly-shaped dimer, where it interacts with membrane lipids [39,40]. Through self-association, VP40 undergoes structural rearrangements into hexameric and higher-order oligomeric assemblies, which are essential for driving membrane curvature and forming the matrix layer beneath the viral envelope [39–41]. This transition is crucial for virion budding, facilitating the release of newly formed viral particles into the extracellular environment.

Beyond its role in virion assembly and release, VP40 also interacts with other viral components, notably the glycoprotein (GP), to coordinate the production of infectious virions. Emerging evidence suggests that VP40 may contribute to immune evasion by modulating host immune signaling pathways, thereby enhancing viral survival and propagation [20,39,41,42]. Through these multifaceted activities, VP40 plays a central role in viral replication, pathogenicity, and host-virus interactions.

At the molecular level, critical amino acid residues involved in VP40 dimerization and higher-order oligomerization include A55, H61, F108, A113, M116, L117, L203, I237, and M305 [29]. These residues form a hydrophobic network essential for stabilizing the dimer interface, with L117 playing a particularly pivotal role in linking VP40 protomers.

Protein sequence alignment across *Filoviridae* species (MARV, RAVV, MLAV, RESTV, SUDV, BDBV, TAFV, EBOV, LLOV, and XILV) revealed conservation patterns critical for understanding VP40 functionality. Residues A113, M116, L203, I237, and M305 are semi-conserved across all viruses studied, suggesting preservation of the core structural functions necessary for virion assembly. In contrast, residues A55, M117, and M241 are semi-conserved only among viruses of the genera *Ebolavirus*, *Striavirus*, *Cuevavirus*, and *Thamnovirus*, but show variability in *Marburgvirus* and *Dianlovirus*. Finally, residues H61, F108, and I307 are not conserved, indicating possible genus- or species-specific adaptations at these positions. These conservation patterns are illustrated in Figure S2.

Overall, the high degree of conservation within VP40’s dimer interface supports its essential role in the viral replication cycle and highlights VP40 as a promising target for antiviral strategies aimed at disrupting virus assembly and egress.

Viral protein 40 (VP40) is a dynamic structural protein essential for filovirus assembly and budding. Initially adopting a butterfly-shaped dimer configuration at the plasma membrane, VP40 undergoes conformational rearrangements to form hexameric structures critical for viral particle assembly [39,40]. This structural plasticity enables VP40 to drive membrane curvature and facilitate the budding and release of virions [40,41]. Beyond its structural role, VP40 also collaborates with glycoprotein (GP) to promote the production of infectious viral particles and has been implicated in modulating host immune responses to aid viral evasion [21,39,41,42].

Critical residues for VP40 dimerization and structural transitions include A55, H61, F108, A113, M116, L117, L203, I237, and M305 [29]. Protein alignment analysis showed that residues A113, M116, L203, I237, and M305 are semi-conserved across all filoviruses, while residues A55, L117, and M241 are semi-conserved among *Ebolavirus*, *Cuevavirus*, *Striavirus*, and *Thamnovirus* species, but vary in *Marburgvirus* and *Dianlovirus*. Residues H61, F108, and I307 are not conserved. These findings (Figure S2) underscore VP40’s evolutionary adaptability, which may contribute to differences in viral assembly efficiency and host range.

#### Glycoprotein (GP)

The glycoprotein (GP) of the Ebola virus, embedded in the viral membrane, plays a pivotal role in mediating viral entry into host cells. GP is composed of two subunits, GP1 and GP2, which are linked by a disulfide bridge. GP1 is primarily responsible for host receptor binding, anchoring the virus to target cells, while GP2 mediates the subsequent fusion of the viral and host cell membranes, enabling viral genome entry into the cytoplasm [20,43–45]. Given its essential function in the early stages of infection, GP is a key target for therapeutic interventions and vaccine development.

An active site located within the GP1 subunit, identified by Zhao et al. (2018) [31], plays a critical role in binding small-molecule inhibitors such as imipramine and thioridazine. This site is characterized by the presence of several charged or hydrophilic residues, notably R64, E100, T519, and D522, which facilitate inhibitor binding. Additional residues contributing to the inhibitor-binding interface include L186, L515, M548, L558, I38, L184, L43, Y517, L554, V66, and A101. These residues form a network of interactions crucial for small-molecule recognition and inhibition of viral entry.

A comprehensive sequence alignment of GP proteins from 12 *Filoviridae* species— including MARV, RAVV, MLAV, RESTV, SUDV, BDBV, TAFV, EBOV, BOMV, LLOV, HUJV, and XILV—revealed notable conservation patterns. Residues L43, E100, A101, and Y517 were found to be non-conserved among the viruses, indicating variability in the active site that could affect inhibitor binding across different filovirus species. Residues I38, V66, L184, L186, and L554 are semi-conserved, suggesting some structural retention of binding features despite sequence variation. Importantly, residues R64, L515, T519, M548, and L558 are semi-conserved in most viruses but show a loss of conservation in LLOV, XILV, and HUJV, three more divergent members of the *Filoviridae* family.

These findings, illustrated in Figure S3, highlight the challenge of developing broad-spectrum GP-targeted antivirals, as variability in key binding residues could limit cross-species efficacy. Nevertheless, the identification of semi-conserved residues provides a strategic foundation for designing inhibitors that retain activity against the major human pathogenic filoviruses.

#### Viral Protein 30 (VP30)

A defining feature of filoviruses is the presence of the RNA synthesis cofactor VP30, a protein uniquely found within this viral family [1,46,47]. VP30 plays a critical regulatory role in Ebola virus replication by acting as both an RNA-binding protein and a transcription activator. It binds to viral RNA and is essential for assembling the RNA replication complex, ultimately facilitating the initiation and regulation of viral RNA synthesis. One of its key functions involves interacting with the Ebola virus nucleoprotein (NP), a process critical for efficient viral transcription. Notably, VP30 harbors a C-terminal globular domain essential for RNA binding and transcriptional activation and also contains a zinc-binding domain necessary for maintaining its structural stability and function [1,2,35].

The interaction between VP30 and NP occurs within the C-terminal globular region of VP30, distinctly away from the VP30 dimer interface. This interaction is predominantly hydrophobic, with limited hydrogen bonding between the proteins. Specifically, the NP residue R612 forms a salt bridge with VP30 residue D202, and NP residue Y611 forms a hydrogen bond with VP30 residue E197 [1,2,35]. Beyond these interactions, additional VP30 residues implicated in NP binding and RNA synthesis include G198, L199, D202, Q203, P205, E209, V207, V210, F222, A225, L226, Q229, W230, S234, M237, and F238.

Sequence alignment analysis of VP30 across filoviruses revealed that residue E197 is fully conserved across all species studied. Several other residues, including L199, D202, Q203, V207, F222, A225, L226, W230, M237, and F238, are semi-conserved among the 11 viruses analyzed. The V210 residue shows semi-conservation in viruses of the genera *Ebolavirus* (EBOV, SUDV, TAFV, RESTV, BOMV, BDBV), *Cuevavirus* (LLOV), and *Striavirus* (XILV), but not in *Marburgvirus* (MARV and RAVV) or *Dianlovirus* (MLAV). In contrast, residues E198, P206, and E209 are not conserved among the analyzed species (Figure S4).

These findings emphasize that while certain critical residues of VP30 are well conserved— supporting its fundamental role in viral RNA transcription—other regions demonstrate genus-specific variability. Such conservation patterns suggest that VP30 represents a promising, though potentially genus-focused, target for antiviral drug development against *Ebolavirus* species.

#### Viral Protein 24 (VP24)

The viral protein 24 (VP24) is a protein found in viruses found in Ebola and Marburg viruses. In the case of Marburg, VP24 interacts with the Keap1 protein to activate the cytoprotective antioxidant response pathway [36,37]. Edwards and colleagues, in 2014 [38], conducted a study on the viral protein VP24 and identified that the amino acid region 205-212 (DIEPCCGE) of the Marburg virus and the K-loop, when placed in the context of the VP24 structural scaffold, play a critical role in the interaction between VP24 and Keap1. The authors also noted that the amino acid sequence near the K-loop is not well conserved among filoviral VP24 proteins, meaning that the DIEPCCGE sequence differs among the *Bundibugyo ebolavirus* (BDBV), *Tai Forest ebolavirus* (TAFV), *Zaire ebolavirus* (EBOV), *Sudan ebolavirus* (SUDV), *Marburg marburgvirus* (MARV), and *Ravn virus* (RAVV).

In the alignment of VP24 protein from all viruses, it was observed that there is semi-conservation of the amino acid residue D205 within the 10 viruses in the region of amino acids 205-212. The amino acid sequence in the region 205-209 is semi-conserved among the viruses of the genera *Ebolavirus* (EBOV, BDBV, TAFT, SUDV, BOMV, RESTV) and *Cuevavirus* (LLOV). A different sequence among amino acids 205-209 is semi-conserved for viruses of the *Marburg* genus (MARV and RAVV). However, the MLAV virus does not show conservation of this region with the other viruses. Figure S5 in supplementary material depicts these results.

#### Nucleoprotein (NP)

The nucleoprotein (NP) is a critical component of the Ebola virus replication machinery, forming the ribonucleoprotein complex (RNP) together with the viral RNA genome and other viral proteins [1,4,22]. This complex constitutes the minimal essential unit required for viral genome replication. In addition to encapsidating the viral RNA, NP plays a fundamental role in assembling nucleocapsid-like structures, which are crucial for the virus replication cycle and particle formation [4,51,52]. Consequently, NP is indispensable for the survival, replication, and propagation of Ebola virus [1,22].

Functional mapping of NP, conducted by Watanabe, Noda, and Kawaoka (2006) [1], identified two key regions essential for its activity: (i) the region spanning amino acids 1–450, which mediates NP-NP interactions necessary for RNP assembly, and (ii) the region from amino acids 451–600, which is critical for the formation of nucleocapsid-like structures.

Sequence alignment of NP proteins from multiple *Filoviridae* species (EBOV, SUDV, BDBV, TAFV, RESTV, BOMV, MARV, RAVV, MLAV, LLOV, XILV, and HUJV) revealed a high degree of conservation in the 1–450 amino acid region, particularly between residues 150–350. This strong conservation highlights the evolutionary pressure to maintain NP-NP interactions essential for viral replication across filoviruses. In contrast, the region from 451–600, important for nucleocapsid assembly, displayed fewer conserved or semi-conserved residues, suggesting greater evolutionary flexibility in this domain. These results are illustrated in Figure S6.

The differential conservation between NP functional domains underscores a key distinction: while NP’s role in genome encapsidation is highly conserved and critical across filoviruses, structural variations in nucleocapsid assembly may contribute to differences in replication efficiency and pathogenicity between species.

#### Large (L) Protein

The large (L) protein of filoviruses functions as the catalytic core of the viral replication machinery, containing two essential enzymatic domains: the methyltransferase (MTase) domain and the RNA-dependent RNA polymerase (RdRp) domain [6–8]. To assess the conservation of functional regions, sequence alignments were performed across the full-length L protein, encompassing both enzymatic domains.

The MTase domain mediates crucial co-transcriptional modifications of viral RNA, including 2’-O-methylation, which are necessary for mRNA stability and host immune evasion. The MTase active site is defined by a catalytic tetrad composed of residues K1813, D1924, K1959, and E1996 [53]. These residues form the core of the 2’-O-MTase catalytic mechanism conserved in Mononegavirales.

The RdRp domain is indispensable for viral RNA genome replication and transcription and represents a major antiviral target [54]. Schmidt and Hoenen (2017) [55] characterized the catalytic center of the Ebola virus RdRp, identifying residues G741, D742, N743, and Q744 as essential for polymerase function. Mutations at these residues severely impair viral replication, highlighting their critical functional role.

Alignment of L proteins from 12 *Filoviridae* species (MARV, RAVV, MLAV, RESTV, SUDV, BDBV, TAFV, EBOV, BOMV, LLOV, XILV, and HUJV) revealed remarkable conservation of the catalytic residues in both the MTase (K1813, D1924, K1959, E1996) and RdRp (G741, D742, N743, Q744) domains. These findings are presented in Figures S7 and S8.

The absolute conservation of these catalytic centers across filoviruses underscores the indispensable role of the L protein in the viral life cycle and strongly supports targeting the L protein enzymatic sites for the development of broad-spectrum filovirus antivirals.

#### Filovirus Inhibitors

The recent development of therapeutic antibodies and small-molecule inhibitors that protect non-human primates from Ebola virus (EBOV) represents a major milestone in anti-filovirus research. However, despite these advances, there are still no antiviral drugs approved for human use against filoviruses, highlighting the ongoing need for sustained efforts toward (pan-)filovirus drug development [56].

Currently, no compound has been systematically tested against all—or even the majority— of known filoviruses [57]. As illustrated in Figure 4, each antiviral compound has been tested, on average, against only three out of the twelve filoviruses studied, with individual compounds ranging from evaluations against two to six viral species. Unsurprisingly, EBOV and Marburg virus (MARV) were the most studied, while four viruses—BOMV, LLOV, XILV, and HUJV—had no available binding or inhibition data for any of their proteins. Due to this sparse coverage, it remains unclear whether some compounds may possess broader antiviral activity than currently recognized, simply because they have not been tested across a wider range of filoviruses.

Table 2 compiles data on drug candidates reported in literature that inhibit at least one homologous viral protein discussed in this study, suggesting candidates that could be prioritized for cross-species evaluation. Our analysis revealed numerous examples where a single compound targets homologous proteins across multiple filoviruses (Figure 5). Most notably, three compounds—Toremifene and two members of the E-series—demonstrated inhibitory activity against the glycoprotein (GP) of six different filoviruses, emphasizing the potential for cross-genus antiviral strategies.

**Figure 5:**
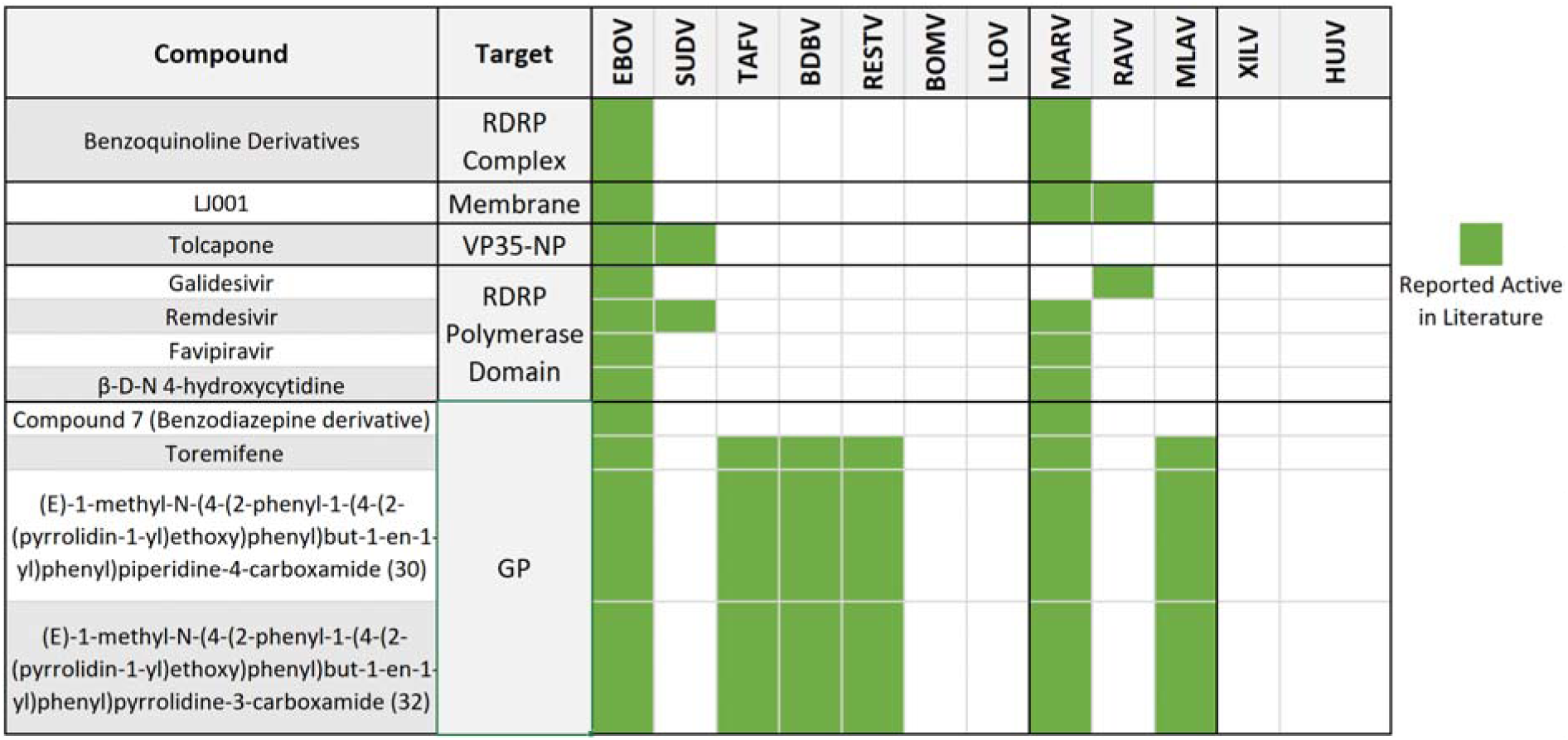
Reported inhibitors that target homologous proteins in multiple filoviruses. VP35-NP refers to the interface of the respective proteins. [58–71]. *Bundibugyo ebolavirus* (BDBV)*, Tai Forest ebolavirus* (TAFV)*, Zaire ebolavirus* (EBOV)*, Sudan ebolavirus* (SUDV)*, Marburg marburgvirus* (MARV and RAVV), *Reston ebolavirus* (RESTV)*, Bombali ebolavirus* (BOMV)*, Lloviu cuevavirus* (LLOV)*, Mengla dianlovirus* (MLOV), and *Xilang striavirus* (XILV).

**Table 2:**
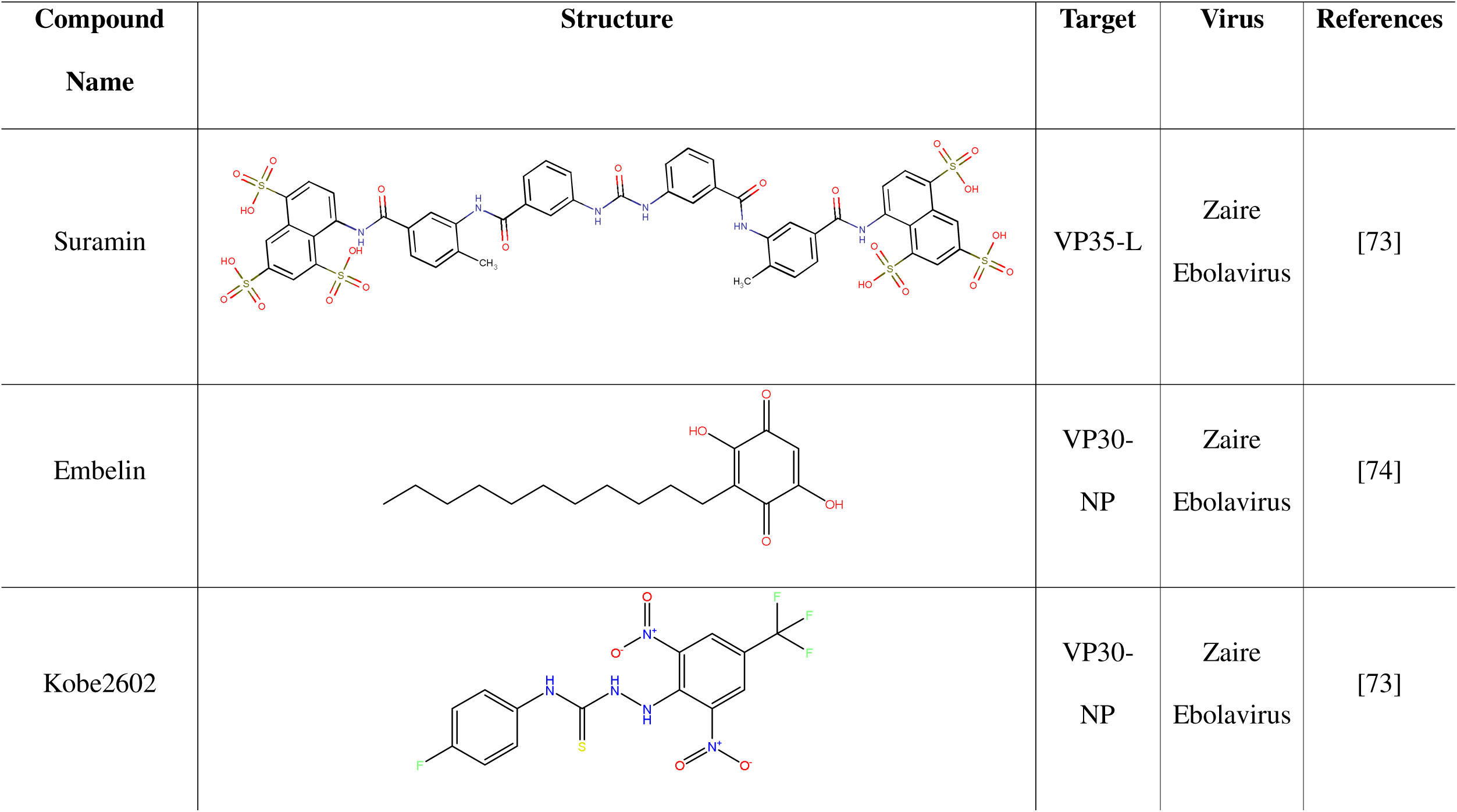
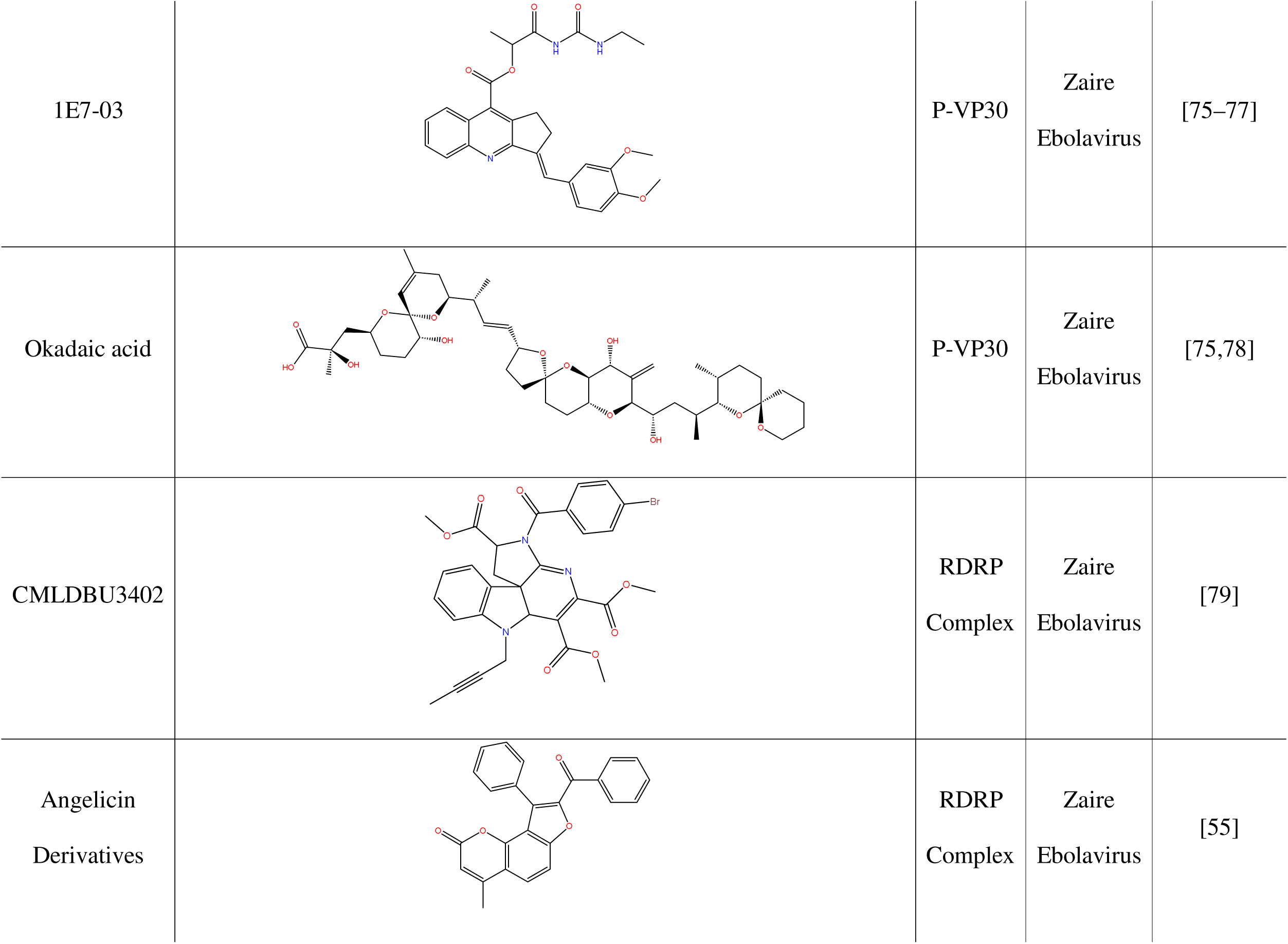
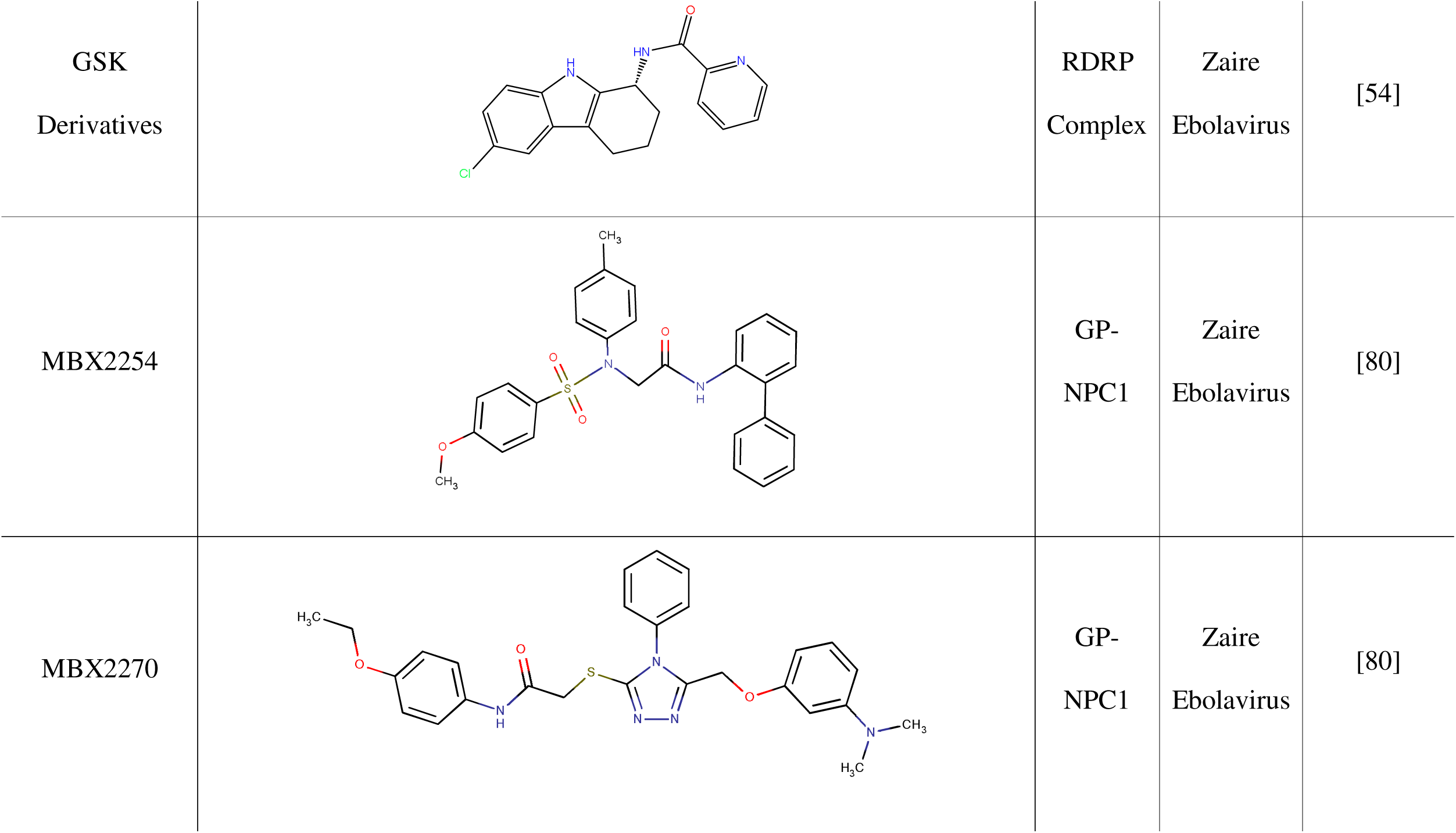
Examples of drug candidates and inhibitors reported in literature [58,76–83]. VP35-L, VP30-NP and GP-NPC1 (Niemann-Pick C1) refer to the respective complexes. P-VP30 refers to the phosphorylated VP30 protein.

| Compound Name | Structure | Target | Virus | References |
| --- | --- | --- | --- | --- |
| Suramin |  | VP35-L | Zaire<br>Ebola virus | [73] |
| Embelin |  | VP30-NP | Zaire<br>Ebola virus | [74] |
| Kobe2602 |  | VP30-NP | Zaire<br>Ebola virus | [73] |
| 1E7-03 |  | P-VP30 | Zaire<br>Ebola virus | [75–77] |
| Okadaic acid |  | P-VP30 | Zaire<br>Ebola virus | [75,78] |
| CMLDBU3402 |  | RDRP<br>Complex | Zaire<br>Ebola virus | [79] |
| Angelicin<br>Derivatives |  | RDRP<br>Complex | Zaire<br>Ebola virus | [55] |
| GSK<br>Derivatives |  | RDRP<br>Complex | Zaire<br>Ebola virus | [54] |
| MBX2254 |  | GP-NPC1 | Zaire<br>Ebola virus | [80] |
| MBX2270 |  | GP-NPC1 | Zaire<br>Ebola virus | [80] |

Focusing on the large (L) protein, the only viral protein with intrinsic enzymatic activity (comprising the RNA-dependent RNA polymerase [RdRp] and methyltransferase [MTase] domains) [6–8], we identified several compounds exhibiting multi-filovirus inhibitory activity. Specifically, Benzoquinoline derivatives were found to inhibit the RdRp complex, while known broad-spectrum antivirals such as Galidesivir, Remdesivir, Favipiravir, and β-D-N4-hydroxycytidine target the RdRp polymerase domain. Notably, Remdesivir, a nucleotide analog prodrug approved by the FDA, exerts its antiviral effect by incorporating into viral RNA and inducing a translocation barrier, thereby stalling the RdRp during elongation [72,73]. Similarly, Favipiravir—a guanidine nucleoside analogue approved in Japan and China—has demonstrated broad-spectrum activity against a variety of RNA viruses, including influenza, SARS-CoV-2, and Lassa virus [56,74]. These findings reinforce the notion that nucleoside analogues continue to represent a highly promising direction for broad-spectrum anti-filovirus drug development.

Our structural conservation analysis above revealed that the catalytic residues of the MTase active site (K1813, D1924, K1959, and E1996) are fully conserved across all twelve filoviruses analyzed. Despite this, no MTase-specific inhibitors have been identified for filoviruses. Based on parallels with efforts to target the flavivirus MTase [75], we propose that the filovirus MTase should be actively pursued as a promising broad-spectrum antiviral target. Furthermore, our results highlight the importance of exploring novel therapeutic strategies, including disrupting key protein-protein interactions critical for the assembly and function of the viral replication complex [1,53,54].

The insights gained from this study extend beyond filovirus research and highlight a broader paradigm shift in antiviral discovery: targeting highly conserved viral proteins offers a strategic pathway to develop resilient, broad-spectrum therapeutics capable of countering both current and future emerging pathogens. By focusing on essential enzymatic functions and replication cofactors such as VP35, VP40, and the L protein’s RdRp and MTase domains—regions under strong evolutionary constraint—researchers can design antivirals less vulnerable to resistance development and viral mutation. This conservation-driven strategy not only streamlines the identification of universal targets within a viral family but also aligns with global pandemic preparedness efforts, aiming to create versatile antiviral platforms that can be rapidly adapted against newly emerging viruses. As the world faces increasingly complex viral threats, leveraging structural and sequence conservation across viral proteomes represents a critical advancement toward durable, next-generation antiviral therapies.

## 4. Conclusion

The persistent threat of filovirus outbreaks, exemplified by Ebola and Marburg viruses, highlights the urgent need for effective, broad-spectrum antiviral therapies. While significant advances have been made—particularly the development of antibodies and small-molecule inhibitors—there remains a notable absence of antiviral drugs approved for human use against filoviruses. Our comprehensive analysis of protein sequence and structural conservation across 12 filoviruses provides critical insights into rational antiviral target selection.

Key replication and assembly proteins—VP35, VP40, VP30, NP, and the L protein— exhibited high conservation across filovirus genera, particularly at functional and catalytic sites essential for viral survival. This underscores the vulnerability of these proteins to therapeutic intervention and supports their prioritization in drug discovery pipelines. In contrast, the surface glycoprotein (GP), while an important vaccine and antibody target, displayed greater sequence variability, especially among non-Ebola filoviruses, suggesting that internal viral proteins may be more suitable for small-molecule, broad-spectrum antiviral development.

Importantly, our study highlights the complete conservation of the catalytic residues within the L protein’s RNA-dependent RNA polymerase (RdRp) and methyltransferase (MTase) domains across all species analyzed. Existing broad-spectrum antivirals such as Remdesivir and Favipiravir target the RdRp, reinforcing the validity of this approach, while the MTase domain remains an underexplored but promising antiviral target. Additionally, structural conservation in VP35 and VP40 provides further opportunities to inhibit critical viral processes such as genome replication, immune evasion, and virion assembly.

Despite these promising findings, we identified significant gaps in antiviral screening efforts: most compounds have only been tested against a small subset of known filoviruses. As a result, the true breadth of many compounds’ antiviral activity remains unknown, suggesting an urgent need for systematic cross-species evaluation.

Overall, our results demonstrate that protein conservation analysis offers a powerful strategy for identifying durable and broad-spectrum antiviral targets. By focusing on highly conserved functional regions, future drug development efforts can improve resilience against viral evolution and emerging filovirus threats. Expanding compound screening across a wider range of filoviruses, coupled with structure-guided drug design targeting conserved sites, holds great promise for delivering the next generation of filovirus therapeutics.

## 6. Acknowledgments

Authors from UNC-Chapel Hill were supported by the National Institutes of Health (Grants U19AI171292 and R01GM140154). The content is solely the responsibility of the authors and does not necessarily represent the official views of the National Institutes of Health.

