## supplementary material for "Conserved Filovirus Proteins as Targets of Broad-Spectrum Antivirals"

**Summary**

**Figure S1:** Alignment of VP35 proteins from all filovirus species: *Bundibugyo ebolavirus* (BDBV)*, Tai Forest ebolavirus* (TAFV)*, Zaire ebolavirus* (EBOV)*, Sudan ebolavirus* (SUDV)*, Marburg marburgvirus* (MARV and RAVV), *Reston ebolavirus, Bombali ebolavirus, Lloviu cuevavirus* and *Mengla dianlovirus*. Highlighted in black rectangles are the amino acid residues of the active site.

**Figure S2:** Alignment of VP40 proteins from all filovirus species: *Bundibugyo ebolavirus* (BDBV)*, Tai Forest ebolavirus* (TAFV)*, Zaire ebolavirus* (EBOV)*, Sudan ebolavirus* (SUDV)*, Marburg marburgvirus* (MARV and RAVV), *Reston ebolavirus* (RESTV)*, Bombali ebolavirus* (BOMV)*, Lloviu cuevavirus* (LLOV)*, Mengla dianlovirus* (MLOV), and *Xilang striavirus* (XILV). Highlighted in black rectangles are the amino acid residues of the active site.

**Figure S3:**Alignment of GP proteins from all filovirus species: *Bundibugyo ebolavirus* (BDBV)*, Tai Forest ebolavirus* (TAFV)*, Zaire ebolavirus* (EBOV)*, Sudan ebolavirus* (SUDV)*, Marburg marburgvirus* (MARV and RAVV), *Reston ebolavirus* (RESTV)*, Bombali ebolavirus* (BOMV)*, Lloviu cuevavirus* (LLOV)*, Mengla dianlovirus* (MLOV), *Huangjiao thamnovirus* (HUJV) and *Xilang striavirus* (XILV). Highlighted in black rectangles are the amino acid residues of the active site.

**Figure S4:** Alignment of VP30 proteins from all filovirus species: *Bundibugyo ebolavirus* (BDBV)*, Tai Forest ebolavirus* (TAFV)*, Zaire ebolavirus* (EBOV)*, Sudan ebolavirus* (SUDV)*, Marburg marburgvirus* (MARV and RAVV), *Reston ebolavirus* (RESTV)*, Bombali ebolavirus* (BOMV)*, Lloviu cuevavirus* (LLOV)*, Mengla dianlovirus* (MLOV), *Huangjiao thamnovirus* (HUJV) and *Xilang striavirus* (XILV). Highlighted in black rectangles are the amino acid residues of the active site.

**Figure S5:** Alignment of VP24 proteins from all filovirus species: *Bundibugyo ebolavirus* (BDBV)*, Tai Forest ebolavirus* (TAFV)*, Zaire ebolavirus* (EBOV)*, Sudan ebolavirus* (SUDV)*, Marburg marburgvirus* (MARV and RAVV), *Reston ebolavirus* (RESTV)*, Bombali ebolavirus* (BOMV)*, Lloviu cuevavirus* (LLOV) and *Mengla dianlovirus* (MLOV). In green, the sequence preserved in the MARV and RAVV viruses, and in yellow, the sequence preserved for the BDBV, TAFV, EBOV, SUDV, RESTV, BOMV and LLOV viruses.

**Figure S6:** Alignment of NP proteins from all filovirus species: *Bundibugyo ebolavirus* (BDBV)*, Tai Forest ebolavirus* (TAFV)*, Zaire ebolavirus* (EBOV)*, Sudan ebolavirus* (SUDV)*, Marburg marburgvirus* (MARV and RAVV), *Reston ebolavirus* (RESTV)*, Bombali ebolavirus* (BOMV)*, Lloviu cuevavirus* (LLOV)*, Mengla dianlovirus* (MLOV), *Huangjiao thamnovirus* (HUJV) and *Xilang striavirus* (XILV). Highlighted in black rectangles are the amino acid residues of the active site.

**Figure S7:** Alignment of MTase/RdRp proteins from all filovirus species: *Bundibugyo ebolavirus* (BDBV)*, Tai Forest ebolavirus* (TAFV)*, Zaire ebolavirus* (EBOV)*, Sudan ebolavirus* (SUDV)*, Marburg marburgvirus* (MARV and RAVV), *Reston ebolavirus* (RESTV)*, Bombali ebolavirus* (BOMV)*, Lloviu cuevavirus* (LLOV)*, Mengla dianlovirus* (MLOV), *Huangjiao thamnovirus* (HUJV) and *Xilang striavirus* (XILV). Highlighted in black rectangles are the amino acid residues of the active site.

**Figure S8:** Alignment of the domain RdRp of L protein from all filovirus species: Bundibugyo ebolavirus (BDBV), Tai Forest ebolavirus (TAFV), Zaire ebolavirus (EBOV), Sudan ebolavirus (SUDV), Marburg marburgvirus (MARV and RAVV), Reston ebolavirus (RESTV), Bombali ebolavirus (BOMV), Lloviu cuevavirus (LLOV), Mengla dianlovirus (MLOV), Huangjiao thamnovirus (HUJV) and Xilang striavirus (XILV). Highlighted in black rectangles are the amino acid residues of the active site.

**Figure S9**: (A) Color-coded depiction of residue conservation at the binding site of all viruses Filovirus NP homologous proteins. Regions red represent conserved residues among homologous proteins. (B) Binding site residues of NP conserved and semi-conserved in all homologous viruses listed in Table 1.

**Figure S10**: (A) Color-coded depiction of residue conservation at the binding site of all viruses Filovirus VP35 homologous proteins. Regions red represent conserved residues among homologous proteins. (B) Binding site residues of VP35 conserved and semi-conserved in all homologous viruses listed in Table 1.

**Figure S11:** (A) Color-coded depiction of residue conservation at the binding site of all viruses Filovirus GP homologous proteins. Regions red represent conserved residues among homologous proteins. (B) Binding site residues of GP conserved and semi-conserved in all homologous viruses listed in Table 1.

**Figure S12:** (A) Color-coded depiction of residue conservation at the binding site of all viruses Filovirus VP40 homologous proteins. Regions red represent conserved residues among homologous proteins. (B) Binding site residues of VP40 conserved and semi-conserved in all homologous viruses listed in Table 1.

**Figure S13:** (A) Color-coded depiction of residue conservation at the binding site of all viruses Filovirus VP30 homologous proteins. Regions red represent conserved residues among homologous proteins. (B) Binding site residues of VP30 conserved and semi-conserved in all homologous viruses listed in Table 1.

**Figure S14:** (A) Color-coded depiction of residue conservation at the binding site of all viruses Filovirus VP24 homologous proteins. Regions red represent conserved residues among homologous proteins. (B) Binding site residues of VP24 conserved and semi-conserved in all homologous viruses listed in Table 1.

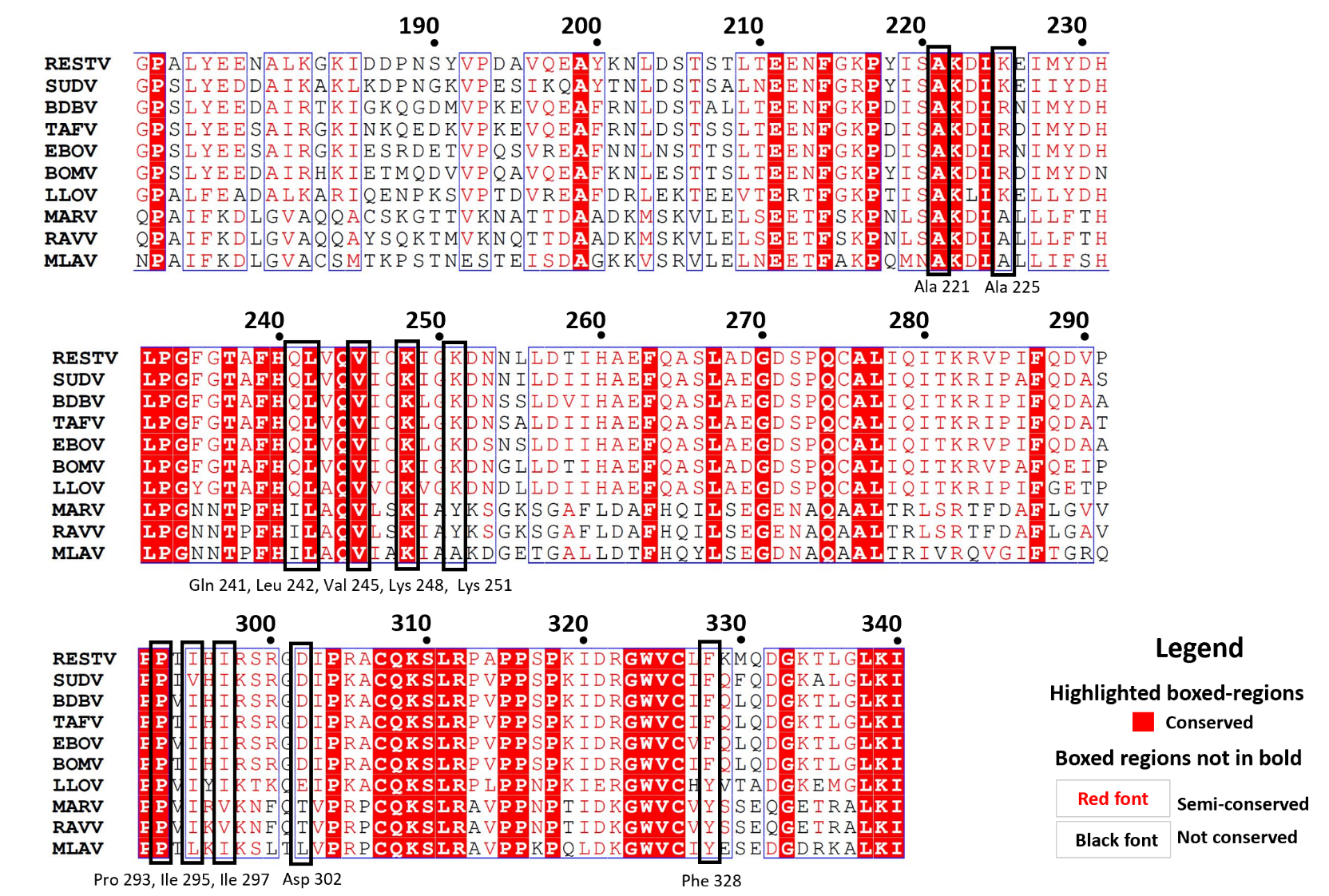

**Figure S1:** Alignment of VP35 proteins from all filovirus species: *Bundibugyo ebolavirus* (BDBV)*, Tai Forest ebolavirus* (TAFV)*, Zaire ebolavirus* (EBOV)*, Sudan ebolavirus* (SUDV)*, Marburg marburgvirus* (MARV and RAVV), *Reston ebolavirus, Bombali ebolavirus, Lloviu cuevavirus* and *Mengla dianlovirus*. Highlighted in black rectangles are the amino acid residues of the active site.

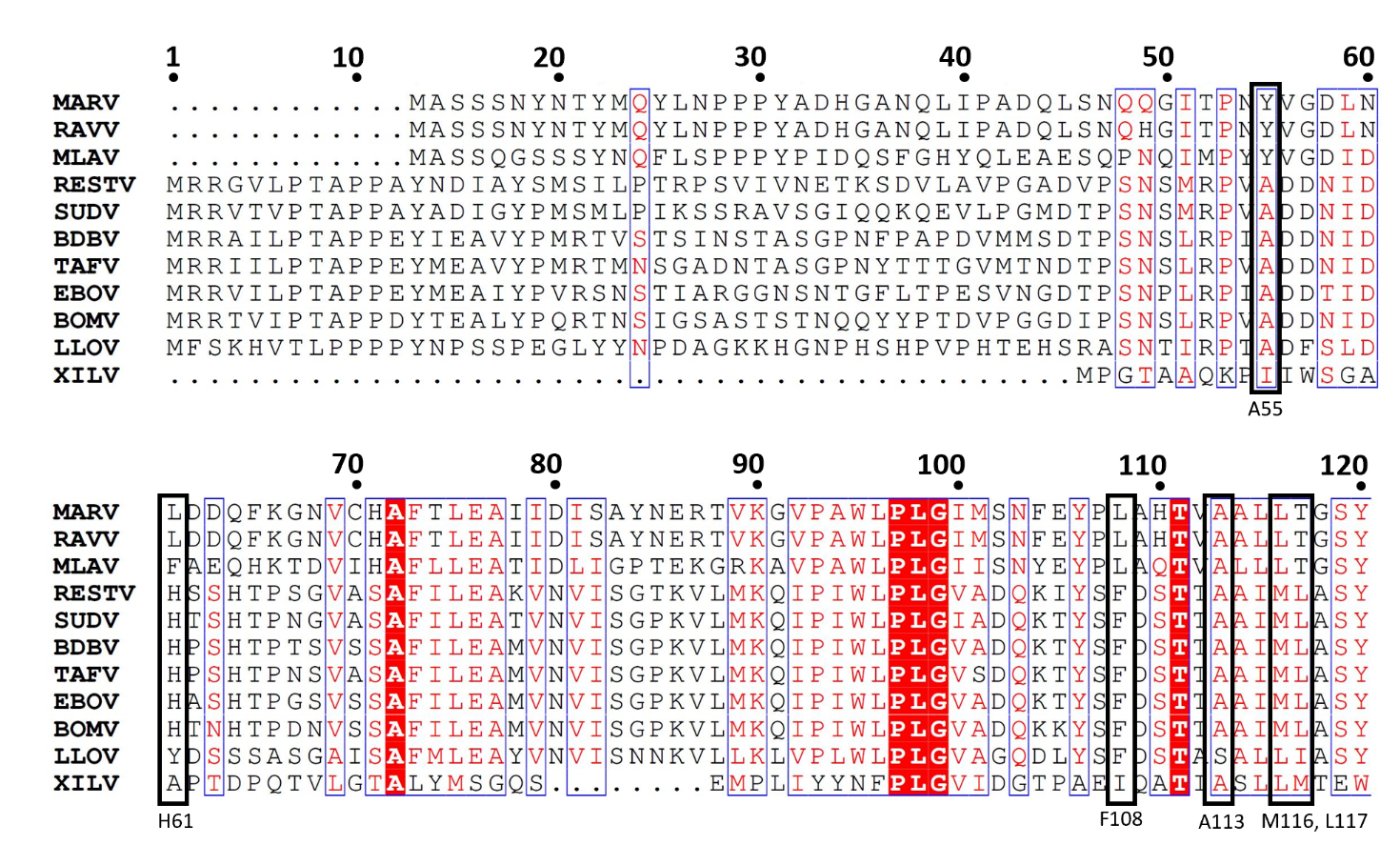

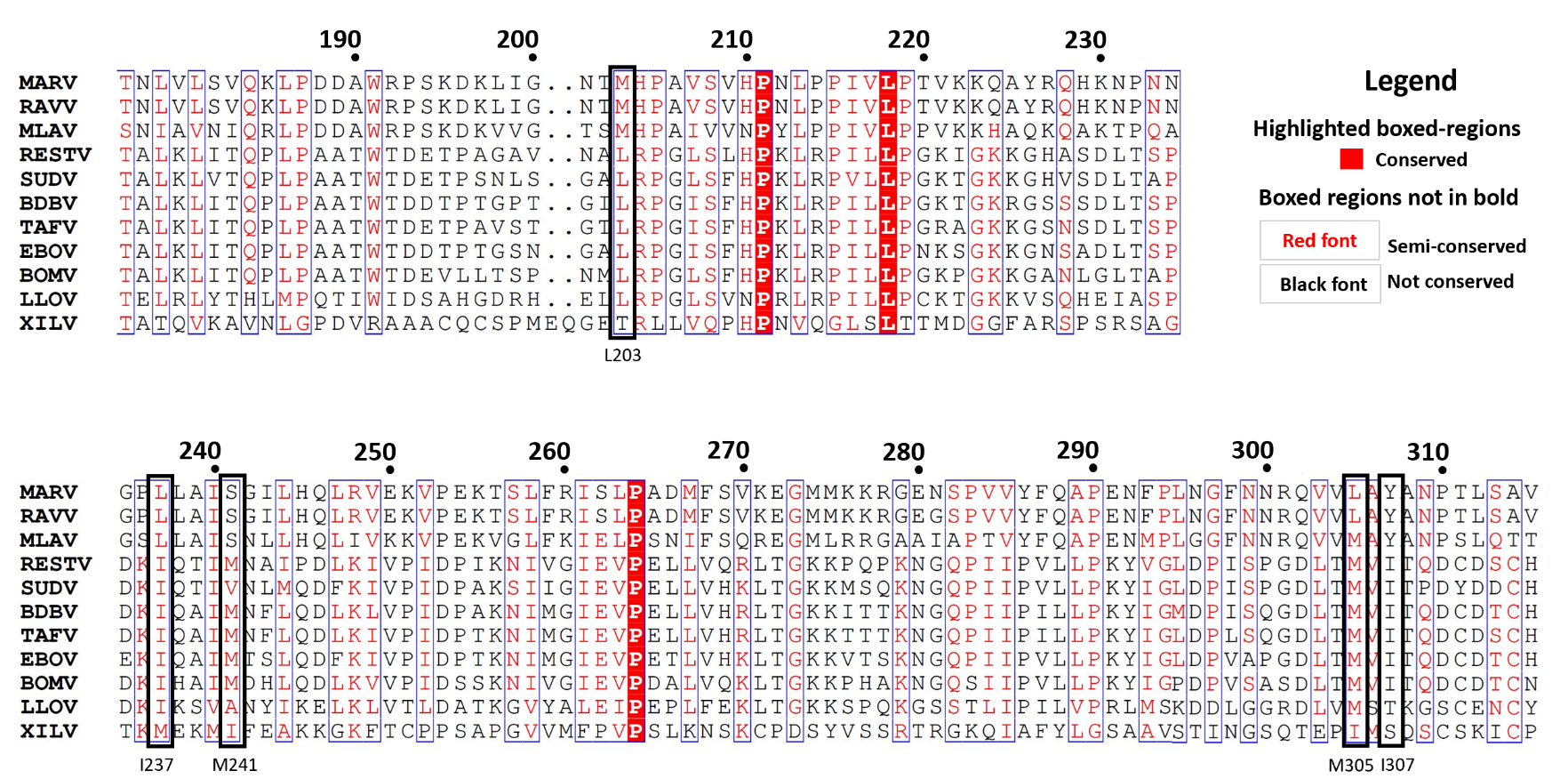

**Figure S2:** Alignment of VP40 proteins from all filovirus species: *Bundibugyo ebolavirus* (BDBV)*, Tai Forest ebolavirus* (TAFV)*, Zaire ebolavirus* (EBOV)*, Sudan ebolavirus* (SUDV)*, Marburg marburgvirus* (MARV and RAVV), *Reston ebolavirus* (RESTV)*, Bombali ebolavirus* (BOMV)*, Lloviu cuevavirus* (LLOV)*, Mengla dianlovirus* (MLOV), and *Xilang striavirus* (XILV). Highlighted in black rectangles are the amino acid residues of the active site.

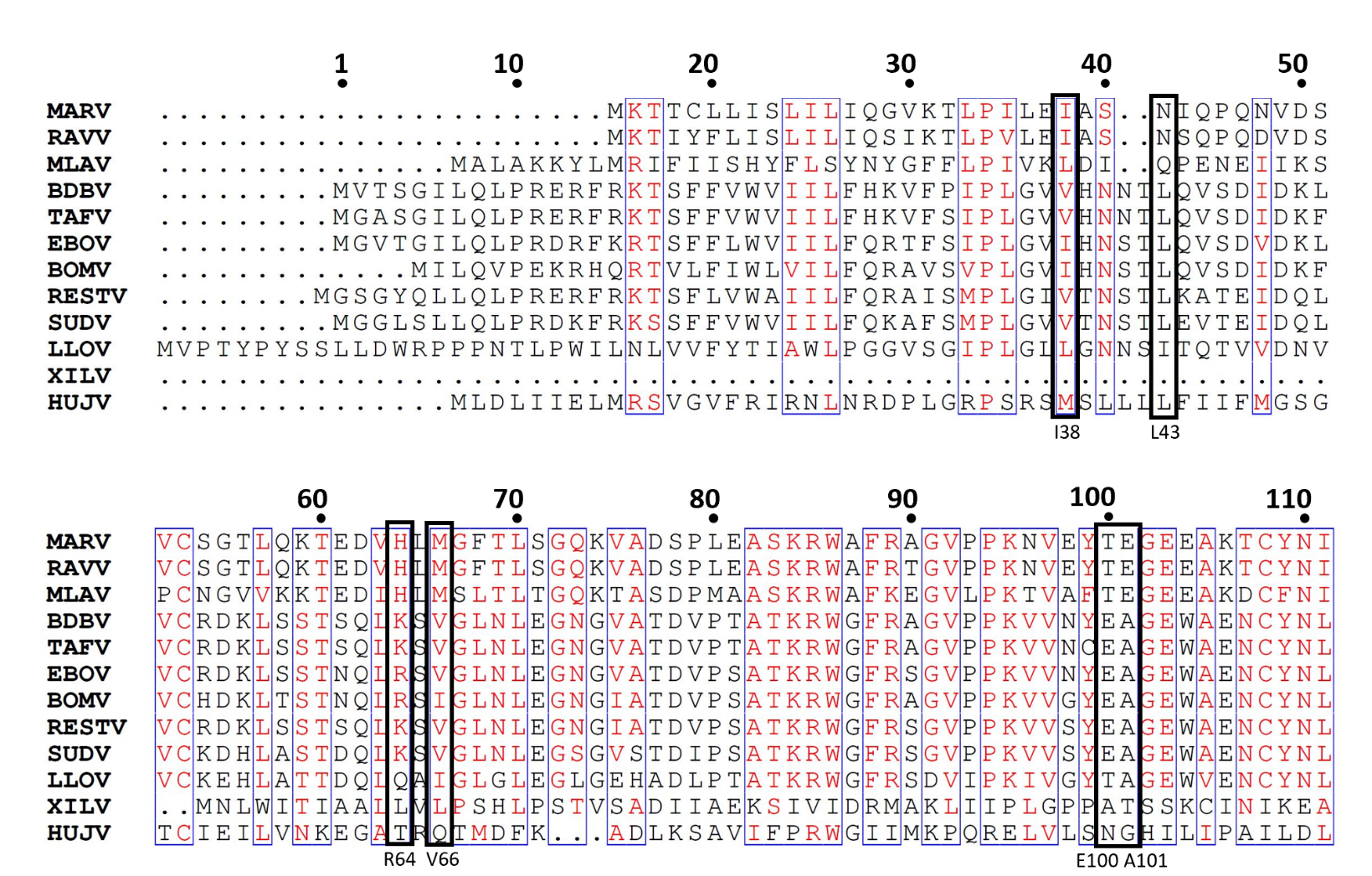

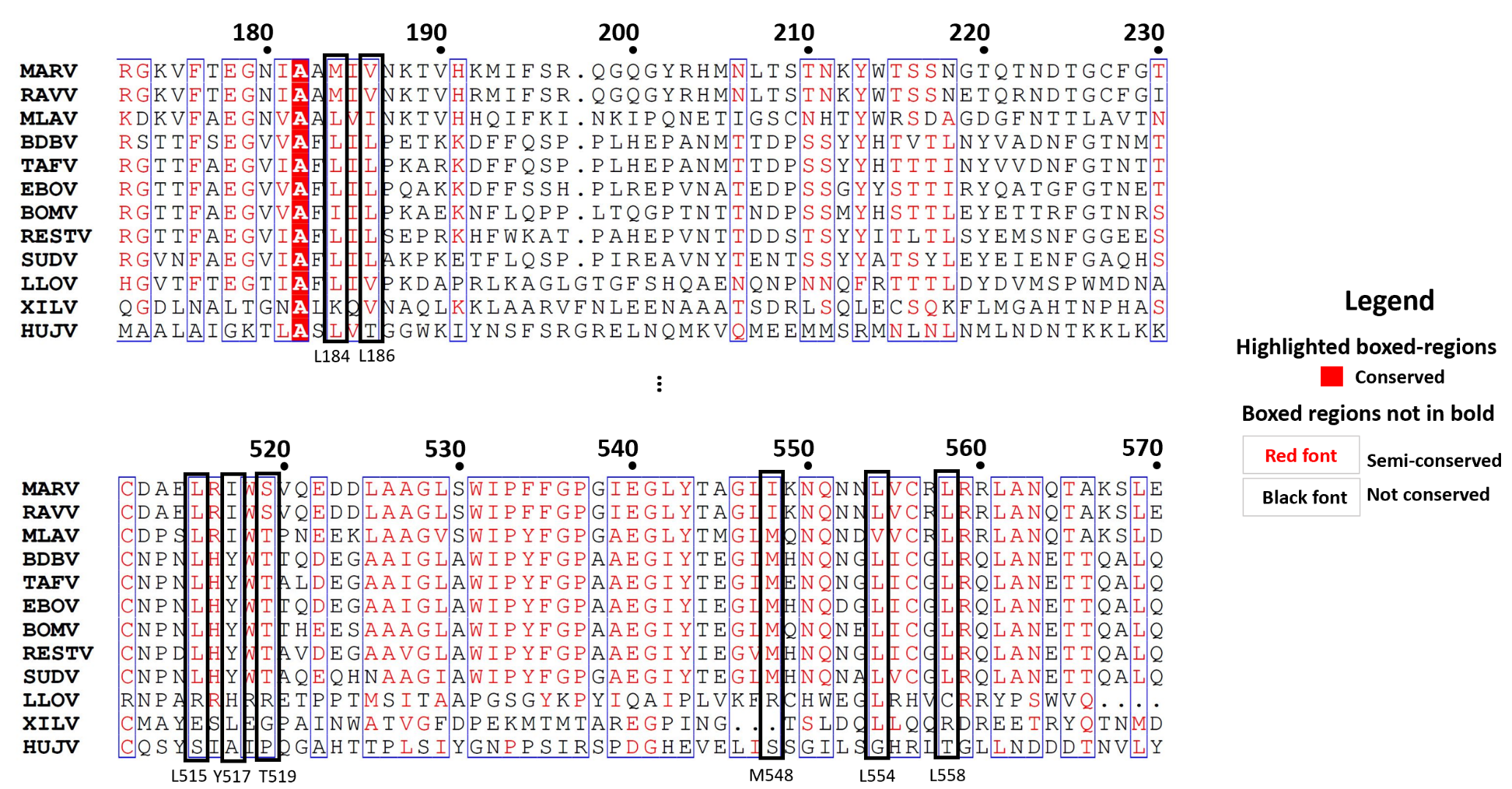

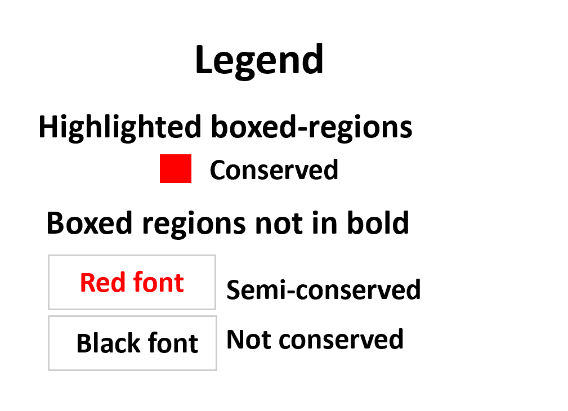

**Figure S3:**Alignment of GP proteins from all filovirus species: *Bundibugyo ebolavirus* (BDBV)*, Tai Forest ebolavirus* (TAFV)*, Zaire ebolavirus* (EBOV)*, Sudan ebolavirus* (SUDV)*, Marburg marburgvirus* (MARV and RAVV), *Reston ebolavirus* (RESTV)*, Bombali ebolavirus* (BOMV)*, Lloviu cuevavirus* (LLOV)*, Mengla dianlovirus* (MLOV), *Huangjiao thamnovirus* (HUJV) and *Xilang striavirus* (XILV). Highlighted in black rectangles are the amino acid residues of the active site.

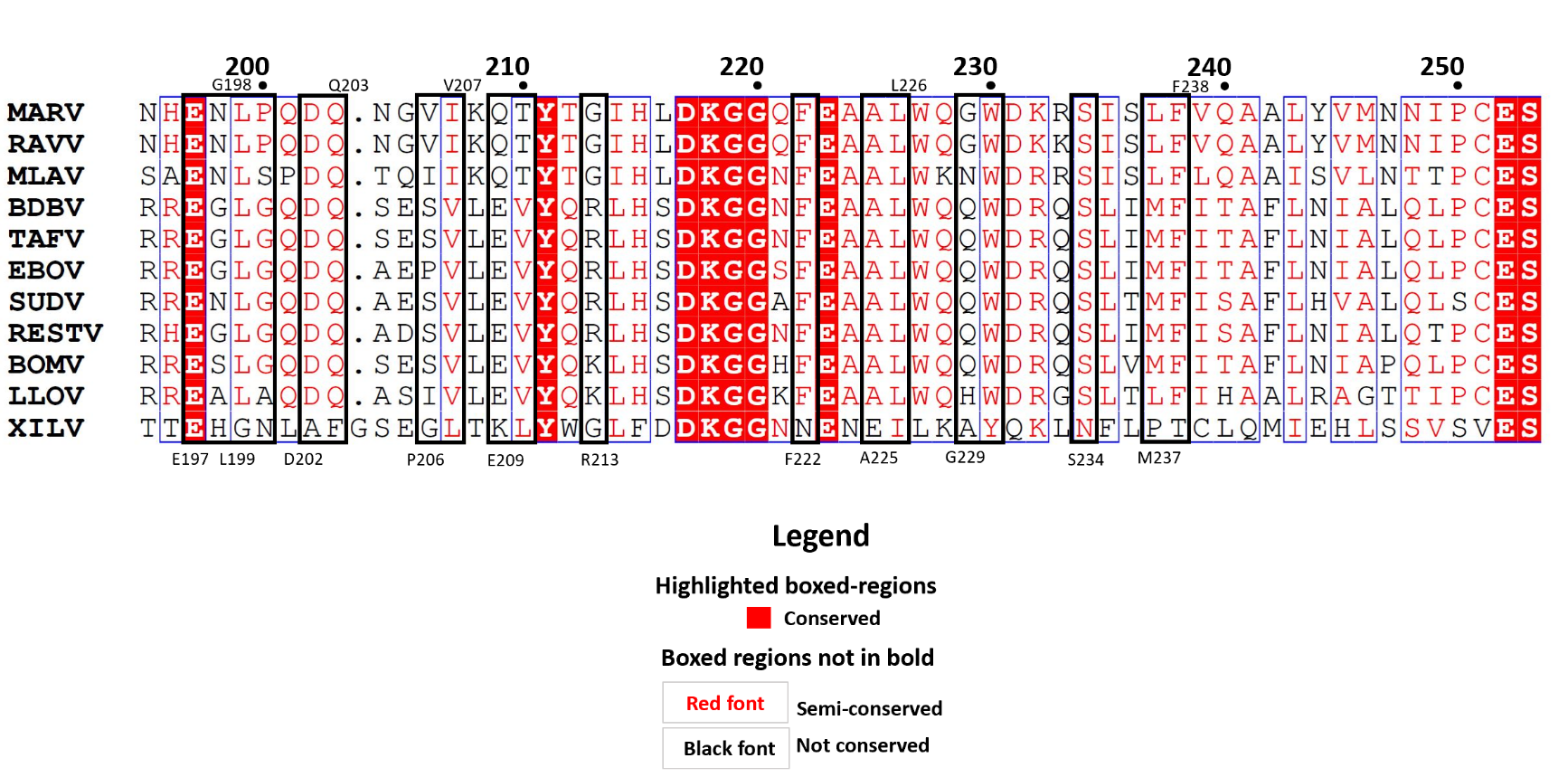

**Figure S4:** Alignment of VP30 proteins from all filovirus species: *Bundibugyo ebolavirus* (BDBV)*, Tai Forest ebolavirus* (TAFV)*, Zaire ebolavirus* (EBOV)*, Sudan ebolavirus* (SUDV)*, Marburg marburgvirus* (MARV and RAVV), *Reston ebolavirus* (RESTV)*, Bombali ebolavirus* (BOMV)*, Lloviu cuevavirus* (LLOV)*, Mengla dianlovirus* (MLOV), *Huangjiao thamnovirus* (HUJV) and *Xilang striavirus* (XILV). Highlighted in black rectangles are the amino acid residues of the active site.

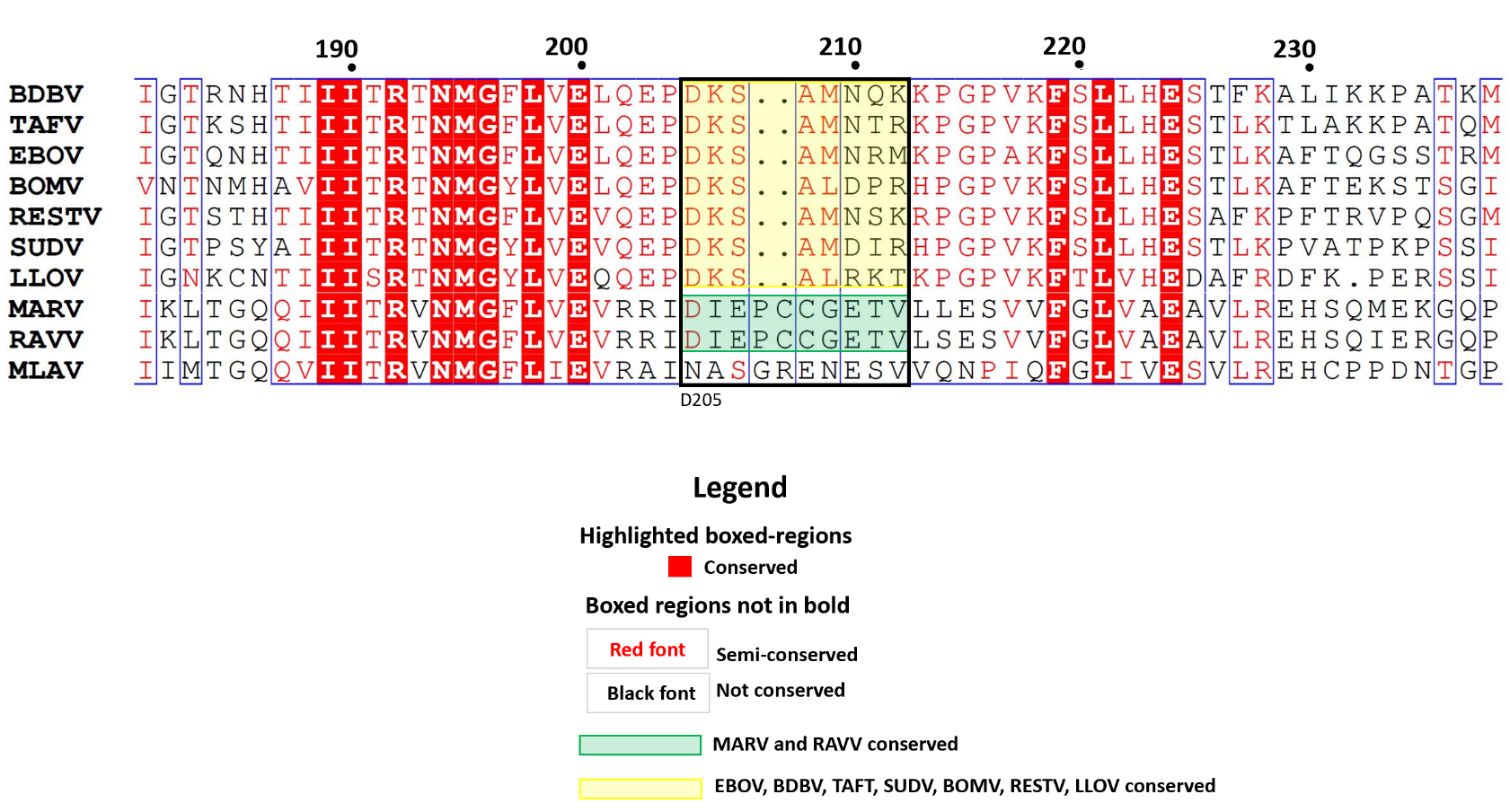

**Figure S5:** Alignment of VP24 proteins from all filovirus species: *Bundibugyo ebolavirus* (BDBV)*, Tai Forest ebolavirus* (TAFV)*, Zaire ebolavirus* (EBOV)*, Sudan ebolavirus* (SUDV)*, Marburg marburgvirus* (MARV and RAVV), *Reston ebolavirus* (RESTV)*, Bombali ebolavirus* (BOMV)*, Lloviu cuevavirus* (LLOV) and *Mengla dianlovirus* (MLOV). In green, the sequence preserved in the MARV and RAVV viruses, and in yellow, the sequence preserved for the BDBV, TAFV, EBOV, SUDV, RESTV, BOMV and LLOV viruses.

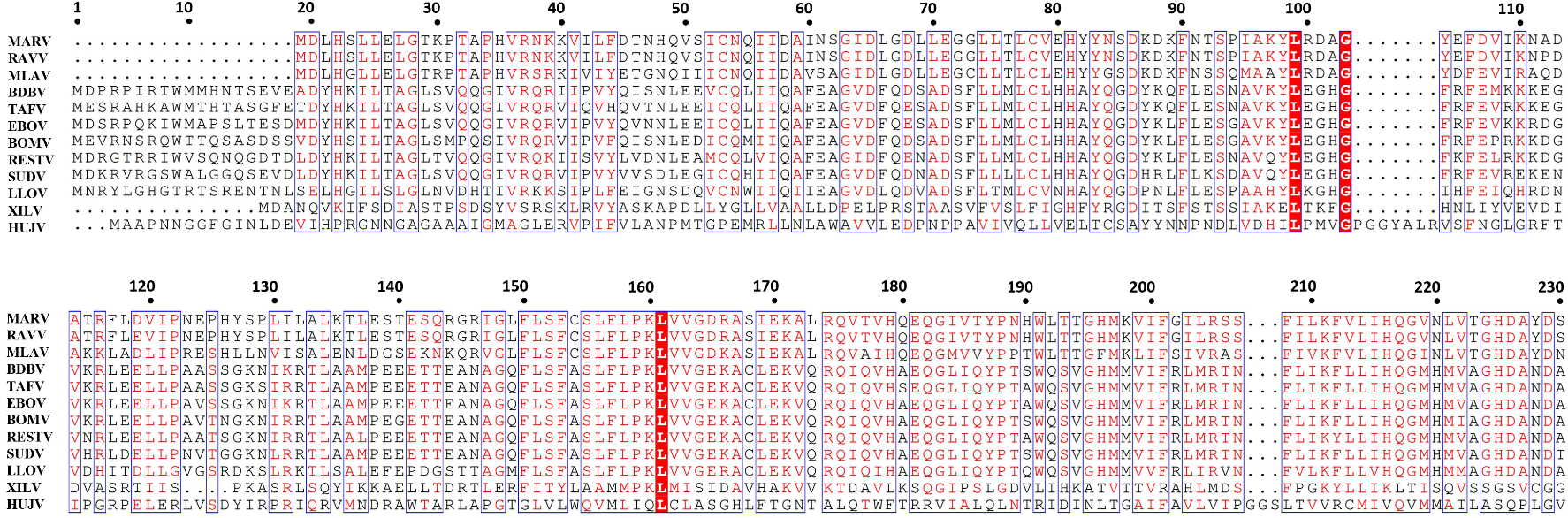

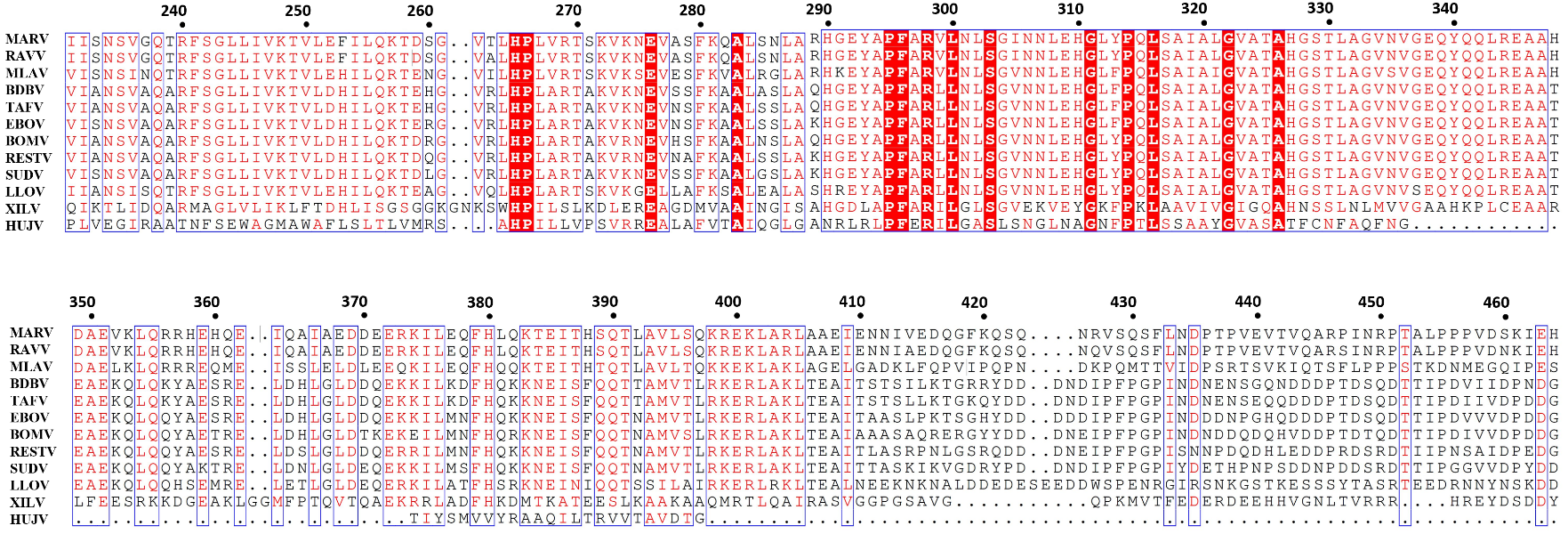

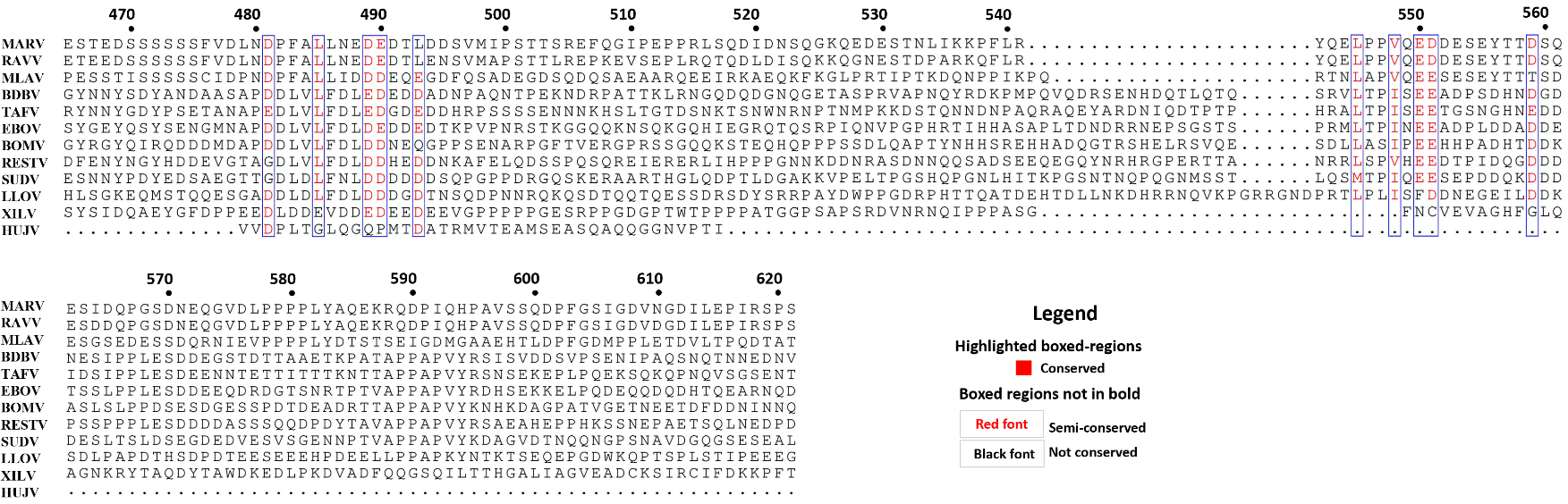

**Figure S6:** Alignment of NP proteins from all filovirus species: *Bundibugyo ebolavirus* (BDBV)*, Tai Forest ebolavirus* (TAFV)*, Zaire ebolavirus* (EBOV)*, Sudan ebolavirus* (SUDV)*, Marburg marburgvirus* (MARV and RAVV), *Reston ebolavirus* (RESTV)*, Bombali ebolavirus* (BOMV)*, Lloviu cuevavirus* (LLOV)*, Mengla dianlovirus* (MLOV), *Huangjiao thamnovirus* (HUJV) and *Xilang striavirus* (XILV). Highlighted in black rectangles are the amino acid residues of the active site.

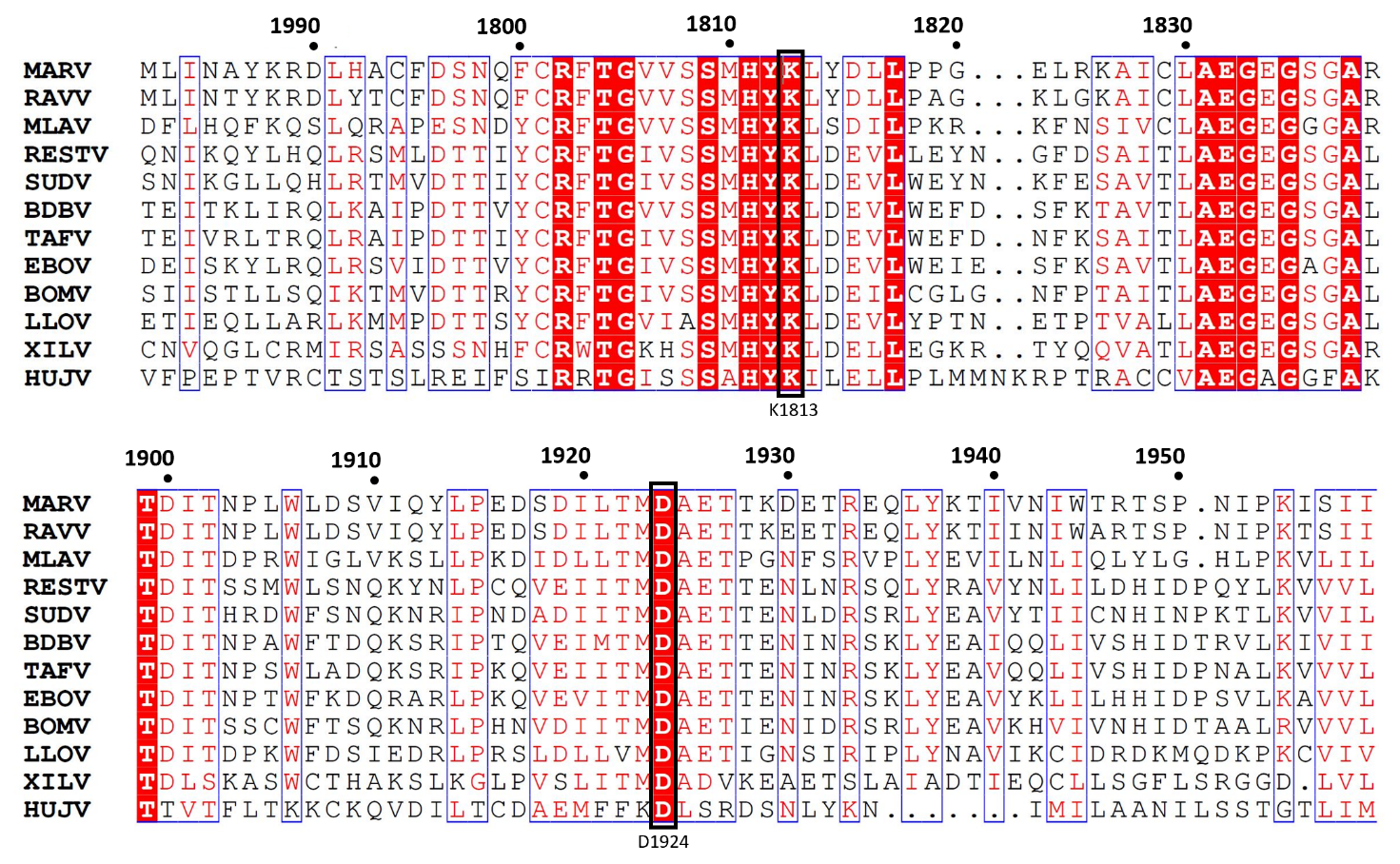

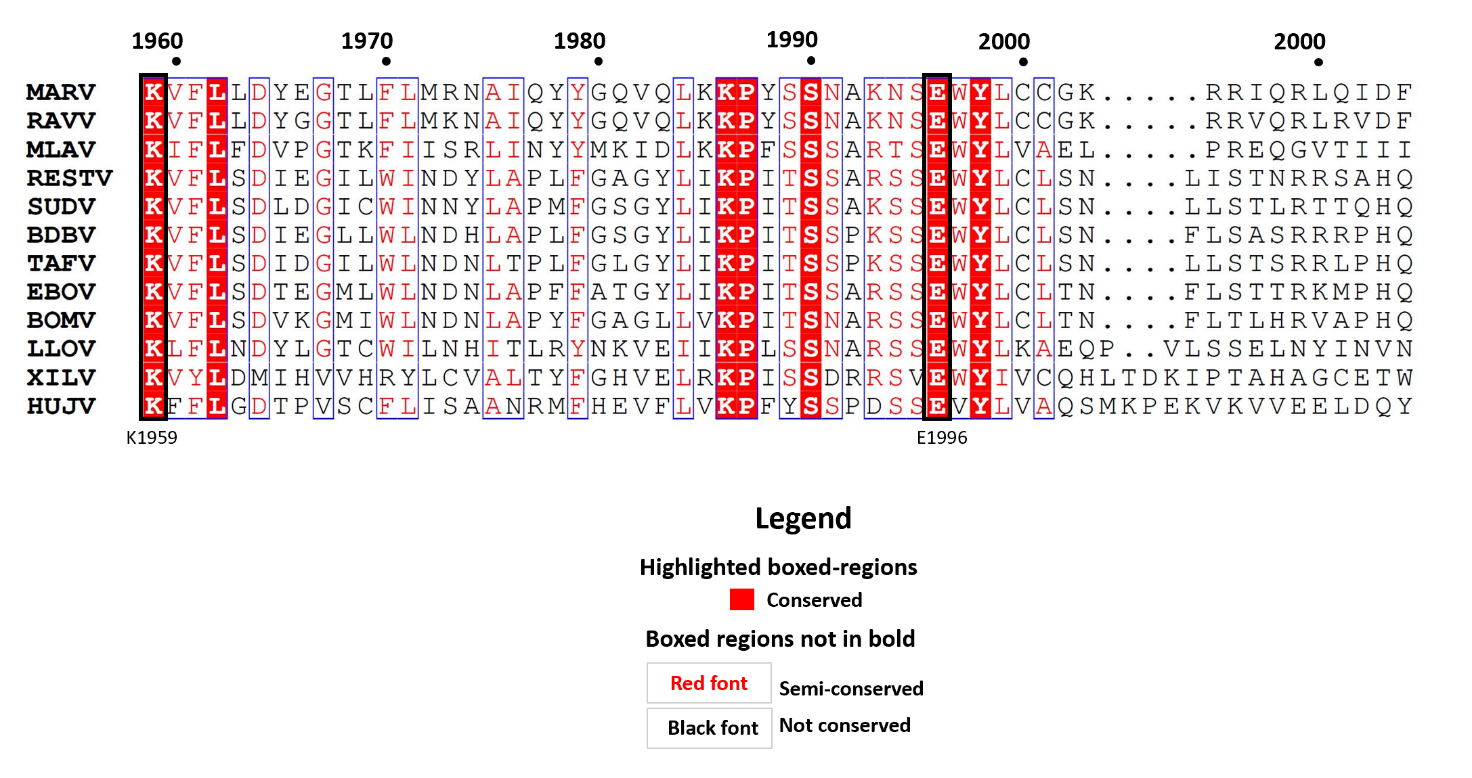

**Figure S7:** Alignment of the domain MTase of L protein from all filovirus species: *Bundibugyo ebolavirus* (BDBV)*, Tai Forest ebolavirus* (TAFV)*, Zaire ebolavirus* (EBOV)*, Sudan ebolavirus* (SUDV)*, Marburg marburgvirus* (MARV and RAVV), *Reston ebolavirus* (RESTV)*, Bombali ebolavirus* (BOMV)*, Lloviu cuevavirus* (LLOV)*, Mengla dianlovirus* (MLOV), *Huangjiao thamnovirus* (HUJV) and *Xilang striavirus* (XILV). Highlighted in black rectangles are the amino acid residues of the active site.

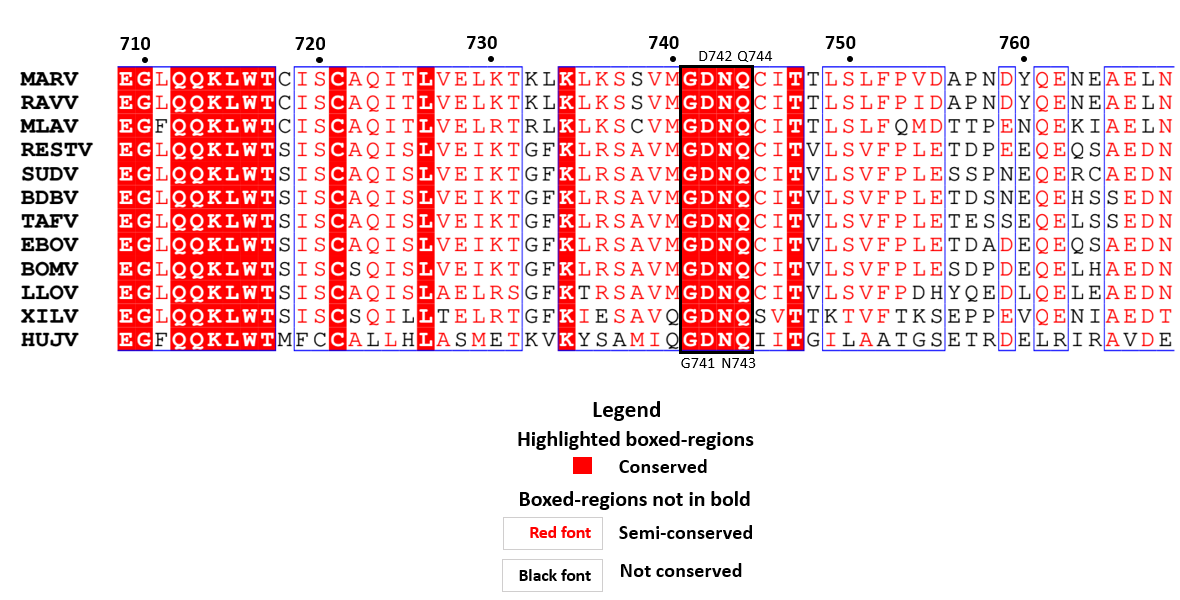

**Figure S8**: Alignment of the domain RdRp of L protein from all filovirus species: Bundibugyo ebolavirus (BDBV), Tai Forest ebolavirus (TAFV), Zaire ebolavirus (EBOV), Sudan ebolavirus (SUDV), Marburg marburgvirus (MARV and RAVV), Reston ebolavirus (RESTV), Bombali ebolavirus (BOMV), Lloviu cuevavirus (LLOV), Mengla dianlovirus (MLOV), Huangjiao thamnovirus (HUJV) and Xilang striavirus (XILV). Highlighted in black rectangles are the amino acid residues of the active site.

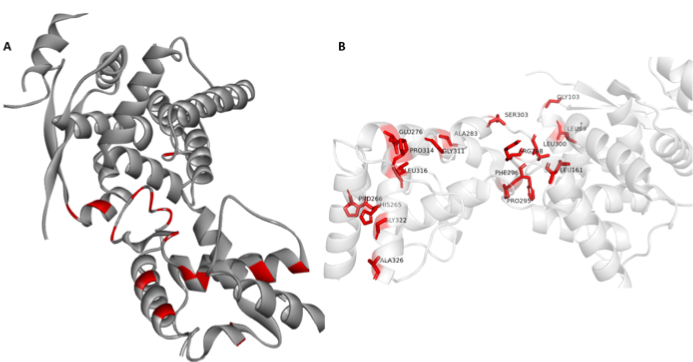

**Figure S9**: (A) Color-coded depiction of residue conservation at the binding site of all viruses Filovirus NP homologous proteins. Regions red represent conserved residues among homologous proteins. (B) Binding site residues of NP conserved and semi-conserved in all homologous viruses listed in Table 1.

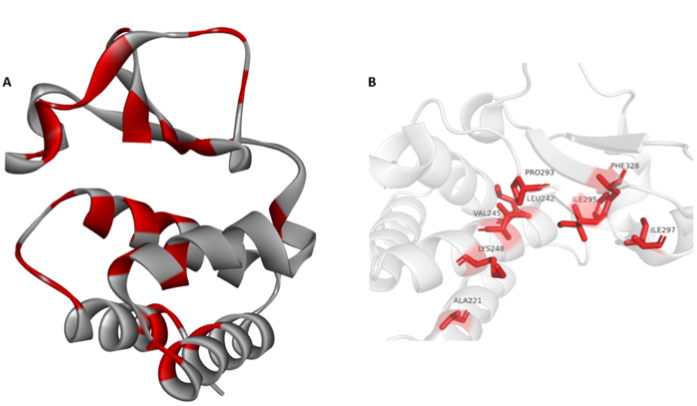

**Figure S10**: (A) Color-coded depiction of residue conservation at the binding site of all viruses Filovirus VP35 homologous proteins. Regions red represent conserved residues among homologous proteins. (B) Binding site residues of VP35 conserved and semi-conserved in all homologous viruses listed in Table 1.

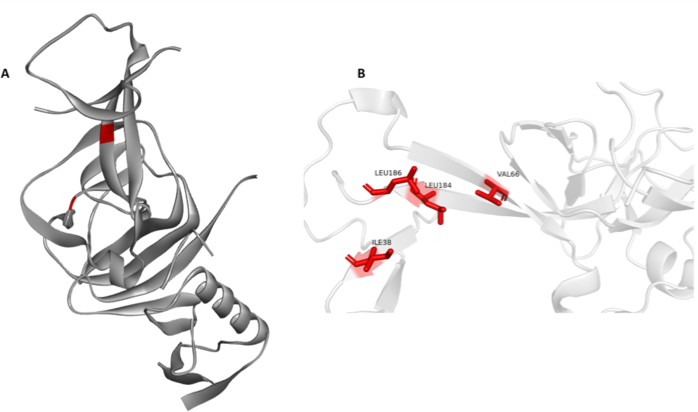

**Figure S11:** (A) Color-coded depiction of residue conservation at the binding site of all viruses Filovirus GP homologous proteins. Regions red represent conserved residues among homologous proteins. (B) Binding site residues of GP conserved and semi-conserved in all homologous viruses listed in Table 1.

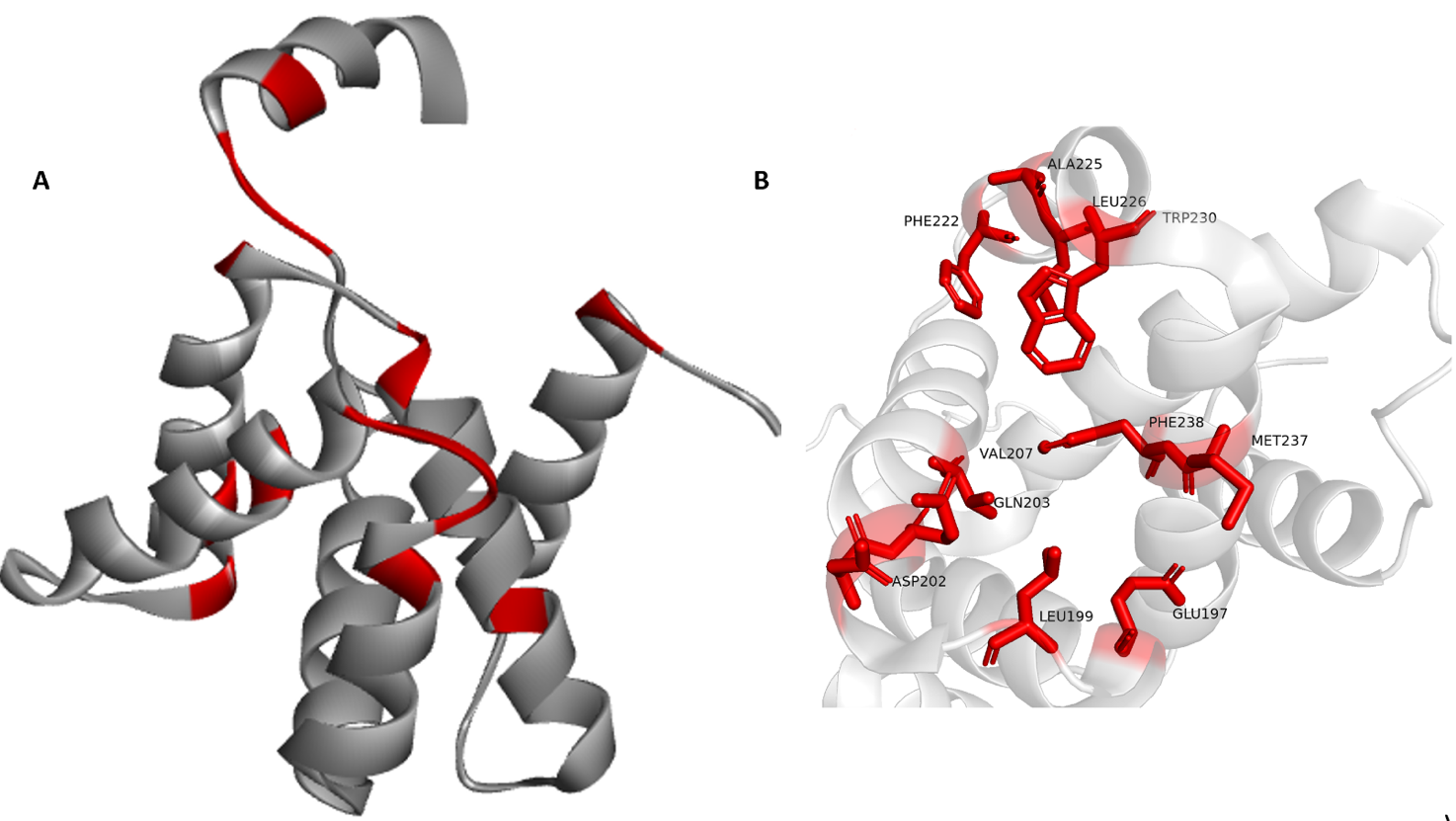

**Figure S12:** (A) Color-coded depiction of residue conservation at the binding site of all viruses Filovirus VP40 homologous proteins. Regions red represent conserved residues among homologous proteins. (B) Binding site residues of VP40 conserved and semi-conserved in all homologous viruses listed in Table 1.

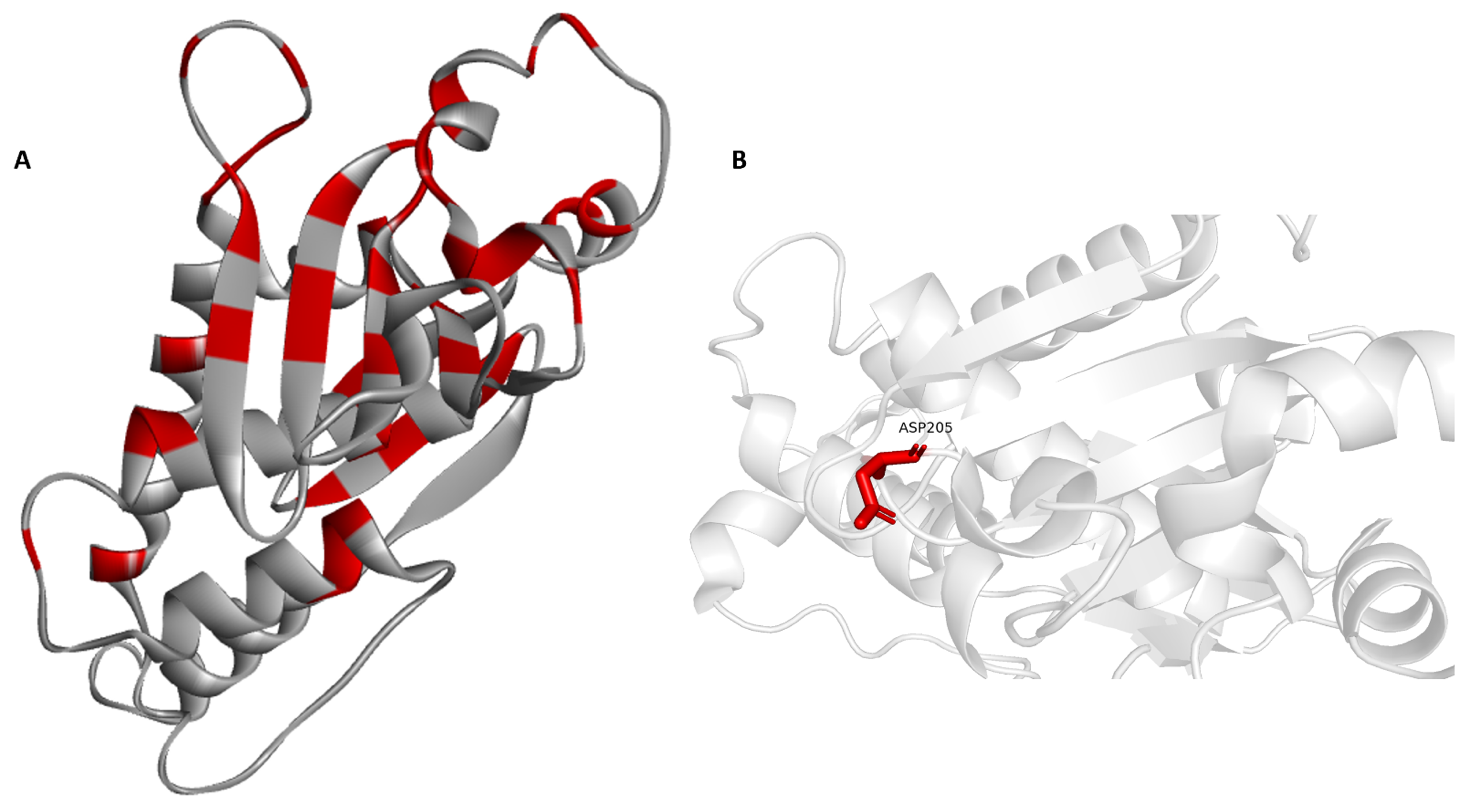

**Figure S13:** (A) Color-coded depiction of residue conservation at the binding site of all viruses Filovirus VP30 homologous proteins. Regions red represent conserved residues among homologous proteins. (B) Binding site residues of VP30 conserved and semi-conserved in all homologous viruses listed in Table 1.

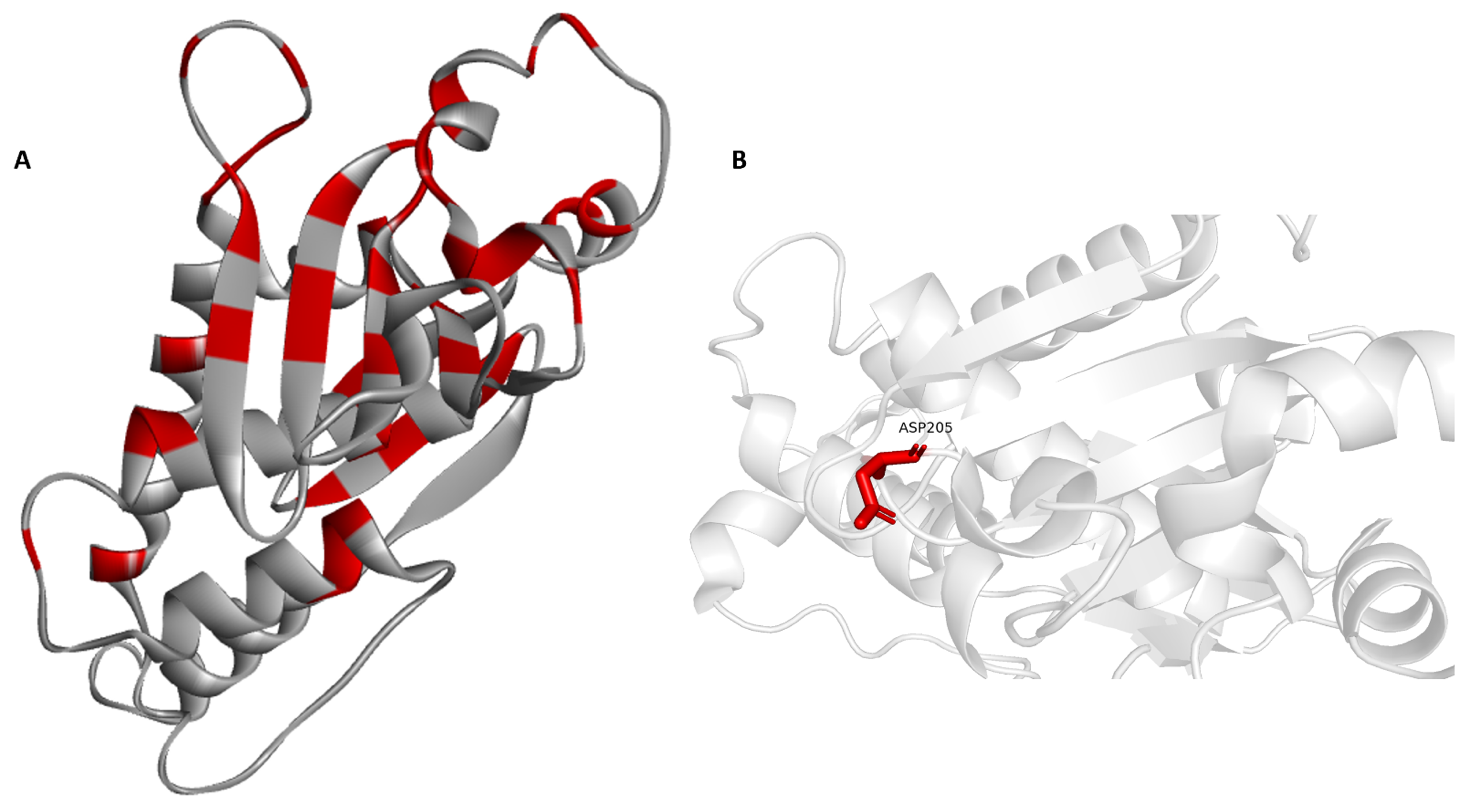

**Figure S14:** (A) Color-coded depiction of residue conservation at the binding site of all viruses Filovirus VP24 homologous proteins. Regions red represent conserved residues among homologous proteins. (B) Binding site residues of VP24 conserved and semi-conserved in all homologous viruses listed in Table 1.
